# FORGEPHAST: A fast, haplotype-first complex population and phenotype simulator for biobank-scale simulation

**DOI:** 10.64898/2026.07.30.741657

**Authors:** Hersh Gupta, Srilakshmi Raj

**Affiliations:** Albert Einstein College of Medicine, Department of Genetics

## Abstract

**Motivation:** Simulation frameworks for statistical and population genetics often separate reference-panel genotype generation, admixture modeling, phenotype simulation, and scalable genomic storage. This limits construction and evaluation of large synthetic cohorts with controlled donor composition, local ancestry, and phenotype architecture.

**Results:** We developed FORGEPHAST, a Python framework for reference-panel-based synthetic haplotype and phenotype generation. FORGEPHAST supports homogeneous mosaic and pulse-admixture generation from user-defined donor panels, integrated genotype quality control, and phenotype simulation with population- and local-ancestry-specific genetic effects. Its HapStore backend provides haplotype-native, Zarr-based storage for scalable numerical analysis. We demonstrate preservation of key donor-panel population-genetic properties and use FORGEPHAST to compare polygenic risk score methods across 54 simulated phenotypic architectures at biobank scales, showing that relative performance depends on phenotype architecture, ancestry, and implementation-specific variant coverage.

**Availability and implementation:** FORGEPHAST is implemented in Python and is available as an open-source package at https://github.com/sriraj-lab/forgephast.

## INTRODUCTION

Genome-wide association studies (GWASs) have become central to understanding how common genetic variation (primarily single nucleotide polymorphisms or SNPs) contributes to phenotypic variability(Abdellaoui *et al*., 2023). GWAS summary statistics can be converted into polygenic risk scores (PRSs), which aggregate many small-effect variants into individual-level risk predictions. For some traits, PRSs now capture risk comparable to monogenic mutations(Kalia *et al*., 2024) and have begun to inform clinical stratification(Lauffer *et al*., 2025). Yet PRSs trained in one ancestry group transfer poorly to others, motivating a growing body of methods aimed at cross-ancestry prediction(Kachuri *et al*., 2024).

Progress on these methods is bottlenecked by data access. Few large-scale biobanks are publicly available, and those that are, restrict individual-level data to secure cloud environments. They also often oversample one ancestry(Corpas *et al*., 2025). Realistic synthetic data is therefore essential for rapid methodological iteration, particularly methods that address multi­ancestry related research questions.

Existing simulation tools fall broadly into two categories: coalescent-based approaches that model ancestral processes directly (e.g., msprime(Baumdicker *et al*., 2022)) and reference-based approaches that generate synthetic haplotypes by mosaic copying from donor panels, typically under a Li–Stephens model and preserve linkage disequilibrium structure(Li and Stephens, 2003). However, current tools have important limitations when applied to diverse, biobank-scale simulation. First, many reference-based simulators were not designed with human population diversity as a primary objective. HapGen2(Su *et al*., 2011) one of the most widely used tools, and G2P(Tang and Liu, 2019) both require substantial manual intervention to simulate multi-ancestry cohorts. HAPNEST(Wharrie *et al*., 2023) targets this gap but restricts users to six predefined continental ancestry labels, limiting flexibility for finer-grained population structure. Second, options to simulate admixed individuals is either absent or handled by separate, smaller-scale tools that lack end-to-end genotype-to-phenotype pipelines for large-scale biobank cohorts(Hou *et al*., 2024; Sun *et al*., 2025). Third, quality control of simulated genotypes and phenotypes is typically left to the user as a post hoc step rather than integrated into the simulation workflow. Fourth, recent empirical evidence suggests that variant effect sizes should be modeled at the local ancestry level, since local haplotype context likely modulates effect magnitudes(Hu *et al*., 2025) — an approach implemented in admix-kit(Hou *et al*., 2024) but not available in most general-purpose simulators. This also necessitates output of phased haplotype data, not just unphased genotypes.

From an engineering standpoint, biobank-scale simulation introduces additional technical demands. Speed is critical for testing many population architectures. Storage formats matter: while PLINK2(Chang *et al*., 2015) provides fast binary haplotype I/O, its custom binary format is not well suited to cloud-native analysis environments. As biobanks increasingly operate within cloud platforms, storage solutions that support both haplotype-oriented access patterns and cloud-native random access will become critical infrastructure. Meanwhile, the performance gap between compiled languages and high-level scientific computing has narrowed considerably, as highly optimized NumPy, JAX, or R routines can match or exceed naive C++ implementations, reducing the advantage of custom compiled code.

To our knowledge, no single existing tool addresses all these requirements. We present the Fast Origin-aware Recombination-driven Genome and Phenotype Amplifier and Simulation Toolkit (FORGEPHAST), a unified framework for diverse, biobank-scale genotype and phenotype simulation. FORGEPHAST operates in three modes. Its primary mode uses the Li–Stephens model to amplify donor populations through recombination, agnostic to any predefined demographic labels. A weighted-donor mode allows closer approximation of modern population mixing. A full admixture mode uses msprime (Baumdicker *et al*., 2022) to simulate ancestry tracts then applies a copying operation from the donor within each tract. Integrated QC suites are provided for each mode.

In addition to simulating genomic information and approximating LD structure, FORGEPHAST includes a complete phenotype simulation module supporting user-specified heritability, polygenicity, and effect-size distributions, along with non-genetic covariates. FORGEPHAST enables realized heritability and effect sizes to vary across populations, which in turn enables simulation of both global and local heterogeneity in genetic architecture, a feature motivated by the effect-size varying by ancestry tract described above.

To address the storage limitations of biobank-scale simulations, we introduce HapStore, a Zarr-based haplotype storage format ideal for efficient and scalable storage of multidimensional data(Miles *et al*., 2020). HapStore supports sample- and variant-level annotation through its metadata layer, provides random access and Pythonic slicing, and integrates naturally with machine learning frameworks. We demonstrate the utility of FORGEPHAST and HapStore through benchmarks against contemporary simulation tools and provide an illustrative application constructing PRSs using various tools to highlight what phenotypic conditions where different models perform well under, as well as the continued gap in transferability. Pre-generated large-scale synthetic datasets in HapStore formats are publicly available for community use at Zenodo (10.5281/zenodo.21679527). FORGEPHAST and HapStore are available at: https://github.com/sriraj-lab/forgephast and https://github.com/sriraj-lab/hapstore.

## METHODS AND ALGORITHMS

### GENOTYPE GENERATION AND QC METHODS

We provide a brief overview of the different modes FORGEPHAST supports for synthetic haplotype generation; more detailed description of the underlying models and related work can be found in Li and Stephens, 2003(Li and Stephens, 2003); Su et al., 2011(Su *et al*., 2011); Gravel, 2012(Gravel, 2012); and Baumdicker et al., 2022(Baumdicker *et al*., 2022). Additional mathematical assumptions and models are described in **Supplementary Methods**, along with mathematical and implementation details of QC metrics. Figure 1 summarizes the genotype and phenotype simulation modes and QC metrics, as well as the overall architecture of the tool.

**Figure 1.**
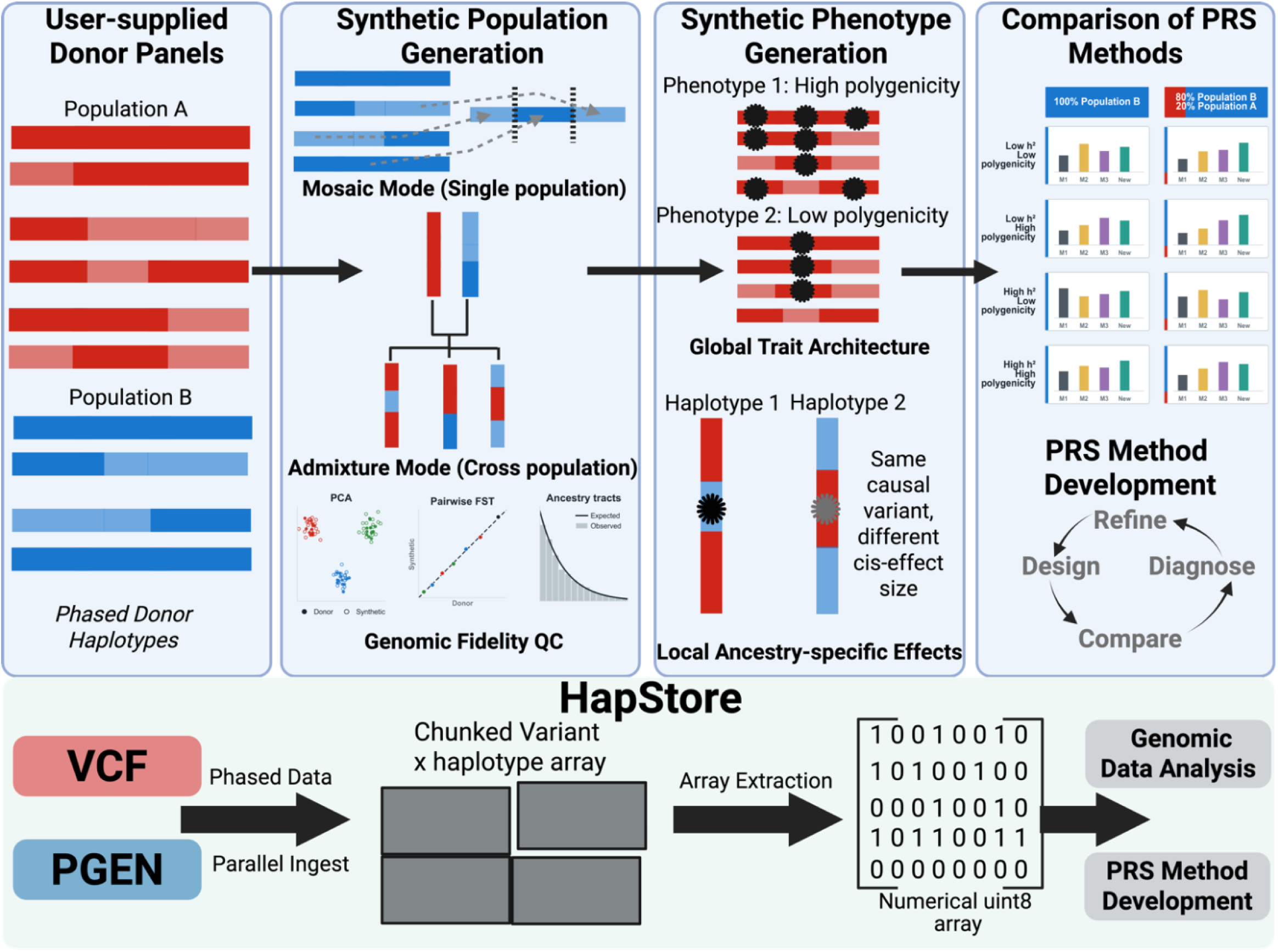
Graphical overview of FORGEPHAST and HapStore. User-supplied phased donor panels are used to generate homogeneous mosaic or admixed synthetic populations with integrated genomic-fidelity QC. Synthetic populations can be paired with controlled global and local-ancestry-specific phenotype architectures for PRS comparison and method development. HapStore provides chunked, Zarr-backed variant-by-haplotype storage with direct extraction into numerical arrays for downstream analysis. Inset plots are illustrative and do not represent quantitative results.

For homogeneous population synthesis, FORGEPHAST uses a Li–Stephens inspired donor-haplotype copying model(Li and Stephens, 2003) in which synthetic haplotypes are formed by concatenating segments copied from a donor panel, with switch probabilities determined by recombination-map distances. Any subset of available donors can serve as the panel, allowing users to define structured subpopulations within larger superpopulations without requiring explicit demographic modeling. An integrated QC suite validates simulation fidelity by comparing donor and synthetic cohorts across multiple axes: pairwise and within-population *F*ST (with automated gating), LD decay curves, allele frequency spectra, KING-robust kinship distributions(Manichaikul *et al*., 2010), alignment with PCA projections, Hardy–Weinberg equilibrium, inbreeding coefficients, and runs of homozygosity. These diagnostics run directly on the simulated output without requiring external tools.

FORGEPHAST also supports a weighted-donor mode, in which donor haplotypes are sampled with user-specified weights rather than sampled from a uniform distribution. While weighted donor sampling has been used elsewhere to approximate admixture, we suggest it is better suited for modeling continuous population structure — cases where subpopulations are closely related and discrete boundaries are artificial. For example, a simulation of South Asian (SAS) individuals from the 1000 Genomes Project might aim to distinguish Northern/Western South Asian and Southern South Asian groups while reflecting the clinal structure observed in practice(Reich *et al*., 2009). A weighted-donor configuration can produce a demographically plausible Northern/Western South Asian cohort by drawing primarily from Northern/Western South Asian donors with a minority contribution from Southern South Asian donors, generating graded structure without requiring a formal admixture model (ancestry tracts, admixture timing, or pulse parameters) and without having to build a new panel for each cline.

For admixed population synthesis, FORGEPHAST uses msprime to simulate ancestry tracts under a discrete-time Wright–Fisher model(Baumdicker *et al*., 2022; Nelson *et al*., 2020), then fills each tract by copying from the appropriate source population’s donor panel. Users specify source populations, their proportional contributions, and an admixture time in generations; msprime produces per-haplotype ancestry tract boundaries, and FORGEPHAST maps these onto the variants and copies donor segments within each tract. This preserves the correlation structure between local ancestry and haplotype content that is critical for benchmarking local-ancestry-aware PRS methods. FORGEPHAST can first generate synthetic source-population panels with the mosaic engine and then use those aligned panels as ancestry-specific donors for admixed simulation. Because admixed haplotypes are copied from the same source-specific HapStore variant axis, the resulting source and admixed cohorts can be analyzed jointly without variant-coordinate harmonization.

An integrated QC module compares the observed distribution of tract lengths and per-individual ancestry proportions against their theoretical expectations under the pulse admixture model(Liang and Nielsen, 2014). Detailed mathematical and implementation details for these QC methods may be found in the **Supplementary Methods**.

### PHENOTYPE GENERATION AND QC

FORGEPHAST simulates continuous and binary phenotypes under a sparse polygenic architecture. A configurable number of causal variants are drawn from MAF-stratified buckets, with hybrid fixed and subset sampling that allows for run-level and replicate-level causal sets. Effect sizes are drawn from configurable base distributions (normal, Laplace, Student-t, or a mixture of normals) and these configurations are modular to allow for other distributions. Effect sizes are rescaled based on MAF(Schoech *et al*., 2019) and can vary across populations through a correlation jitter model, where effects must still correlate with each other and are not allowed to deviate freely. The correlation can either be manually specified or derived from pairwise *F*ST calculations. An optional infinitesimal genetic background component can be added, as well as fixed, random, and hybrid non-genetic covariates. All components are rescaled to user-specified variance fractions (heritability, covariate, residual).

Heritability (*h*^2^) is fixed for an overall cohort. Heritability per-population is induced from the jittered effect sizes and a controlling parameter that decides the possible variant choices based on how similar MAF is- this allows for controlling the coupling of realized heritability between populations. As an example, the “tight” setting only allows for a twofold difference in MAF for a given variant between all populations. For admixed individuals, the genetic component is computed by applying locally-inherited source-population effect sizes according to per-haplotype ancestry at each variant, with no additional rescaling. Realized heritability in admixed samples therefore emerges from the mixture of source-population effect sizes, individual-level ancestry proportions, and local haplotype structure, naturally producing the heritability attenuation observed empirically in admixed cohorts when cross-population effect correlation is imperfect.

FORGEPHAST also supports non-uniform distribution of variants across a genome, where a user can supply a file with annotations across the genome and upweight or downweight the chance of selecting variants within those annotations. The default usage of this method causes variants within protein-coding gene bodies and 100 kb upstream of transcription start sites to be upweighted, . Users can also supply custom BED annotations to mimic the biological reality of more active regulatory and coding regions and gene deserts.

FORGEPHAST uses the continuous phenotype module as the basis of generating binary/case-control phenotypes. The conversion of continuous phenotype to binary phenotype is performed by specifying a prevalence, and then fitting a liability model(Falconer, 1965). The prevalence can be specified at the level of population. For admixed individuals, prevalence is interpolated from source populations through local ancestry exposure at causal loci. Exact mathematical structure for the liability and prevalence models can be found in the **Supplementary Methods**.

For phenotypic QC, FORGEPHAST directly supports running GWAS on the simulated phenotypes. GWAS engines can be flexibly added, but the default shipped engines are currently PLINK2 and REGENIE(Mbatchou *et al*., 2021). FORGEPHAST’s phenotype QC utilizes GWAS validation that compares recovered effect estimates against ground truth and population-weighted betas for admixed samples. An integrated plotting suite produces beta scatter, Manhattan, QQ, causal-variant recovery among top-ranked associations, enrichment, and group-stratified panels. *HAPSTORE STORAGE FORMAT.* To deal with file I/O in a haplotype-native format, we developed a Zarr-based storage format(Miles *et al*., 2020) called HapStore, which stores phased diploid haplotypes as uint8 NumPy arrays. These can be stored using Zstd as compressed, sample and variant chunked array stores, with separate sample and variant metadata. HapStore’s implementation contains separate variant and sample axis merges, allowing operations that only occur on one axis.

HapStore allows for random access of both variants and samples, while allowing for usage of metadata for slicing **(Figure 2A**, **Figure 2B)**. While PLINK2’s PGEN format offers fast sequential scan performance (**Figure 2A**), its binary layout is accessed primarily through PLINK2 itself or a thin C library, and reaching array-oriented frameworks such as NumPy or PyTorch requires an explicit conversion step. HapStore inherits Zarr’s native bindings across Python, R, Julia, and JavaScript, transparent support for cloud object stores, and direct interoperability with scientific computing and machine learning frameworks without custom readers — making it better suited to simulation and analytical workloads where haplotype-level access, flexible metadata, and programmatic slicing are primary requirements.

**Figure 2.**
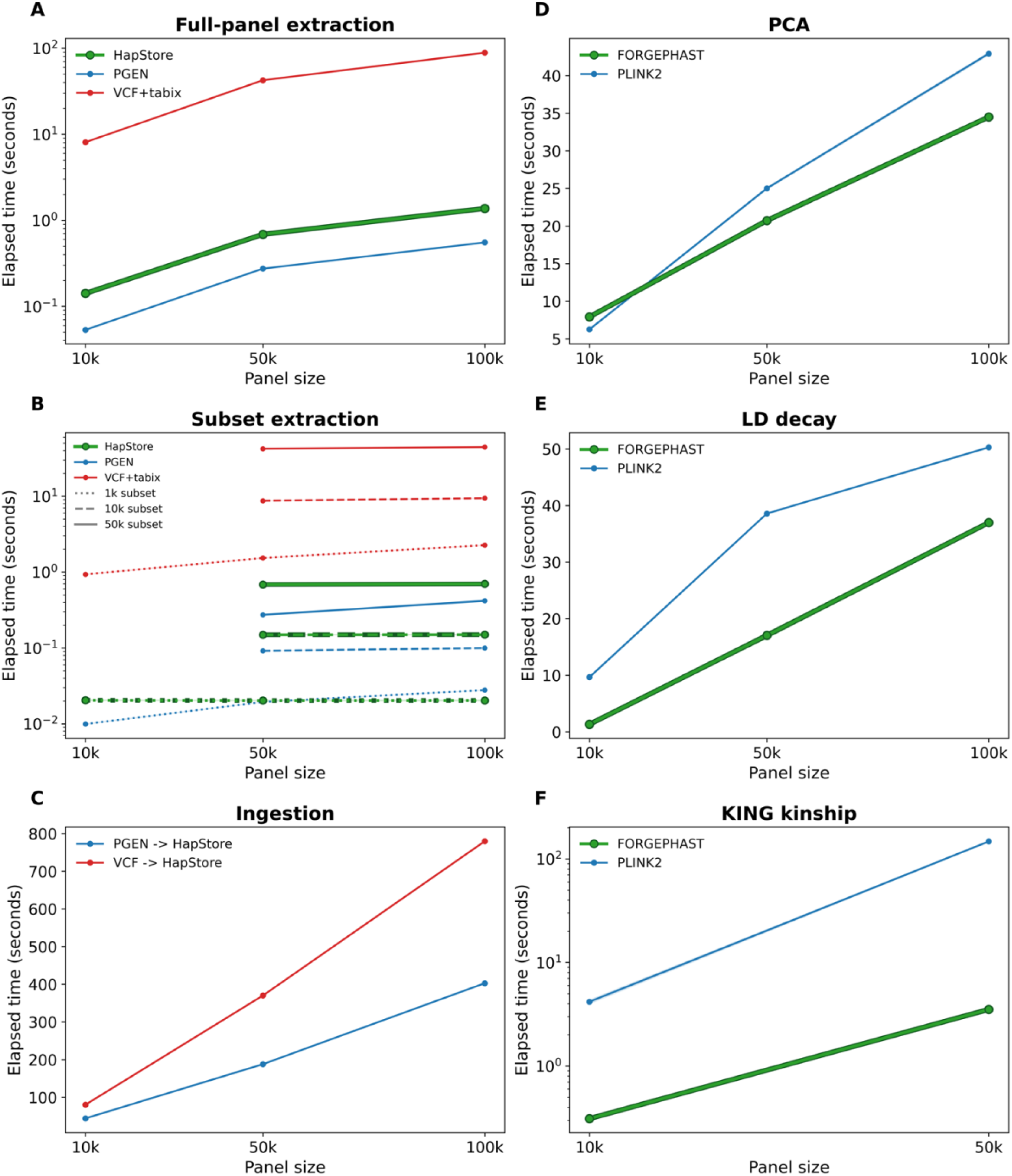
Comparison of HapStore with common genetic data formats and analysis backends. **A:** Time (seconds) to extract a 0.5 cM chromosome 5 window across all samples into an in-memory analysis array. **B:** Time (seconds) to extract 1,000-, 10,000-, and 50,000-sample subsets from 10,000-, 50,000-, and 100,000-sample parent panels, over the same 0.5 cM window into an in-memory analysis array. **C:** Time to convert chromosome 5 files containing 482,118 variants from VCF or PGEN into HapStore format using HapStore ingestion functions. **D-F:** Runtime comparisons between PLINK2 C++ implementations and HapStore-backed Python implementations for common statistical-genetic operations. Exact variant counts, sample sizes, benchmark settings, and function-specific parameters are provided in the Supplementary Methods. FORGEPHAST and HapStore stack are bolded for identifiability.

HapStore is optimized for ingestion of the PGEN and VCF data formats (**Figure 2C)**. Finally, with the usage of high-performance and optimized Python code, QC function kernels in FORGEPHAST built on HapStore (PCA, LD correlation, KING score, etc.) can match or exceed equivalent PLINK2 custom C++ implementations (**Figure 2D, 2E, 2F**; numerical metric parity demonstrated in **Supplementary Figure 1** for AFS and KING score).

Unlike VCF Zarr(Czech *et al*., 2025), which provides a general Zarr encoding of the VCF callset model, HapStore is a haplotype-first storage layer designed for repeated numerical operations on phased panels. Conceptually, HapStore is closer to an analysis-oriented PLINK/PGEN replacement than to a VCF replacement: it stores the phased haplotype matrix needed for simulation and downstream numerical workflows rather than the full VCF callset model.

## RESULTS

### SYNTHETIC POPULATION SIMULATION

As an example of homogeneous superpopulation simulation, a donor panel, consisting of 4 non-admixed combined populations representing AFR, CSA, EUR, and EAS (see **Supplementary Table 2** for exact included populations), was created, representing a total of 1748 donors(1000 Genomes Project Consortium *et al*., 2015). 10k synthetic diploids were created for each population for chromosome 1 (570k variants), chromosome 8 (414k), and chromosome 21 (102k variants). The output of the QC metrics from this simulation, including allele frequency spectrum, LD root mean-square error, pairwise donor vs synthetic *F*ST, KING kinship distributions, and the PC plot of PCs 1 and 2 are detailed in **Figure 3**. All metrics show close agreement between donor and synthetic populations. Pairwise population-separation AUCs were near 1.0 in both the donor and donor-projected synthetic cohorts, with minimal absolute differences between the two (**Supplementary Data 1**). Other metrics, including Hardy-Weinberg statistics and runs of homozygosity, can also be found in **Supplementary Data 1**. Because mosaic recombination draws from unrelated donors without selective pressures, synthetic cohorts have less identity-by-descent than real populations, producing closer adherence to Hardy-Weinberg expectations than observed in natural populations, as previously identified(Wharrie *et al*., 2023).

**Figure 3.**
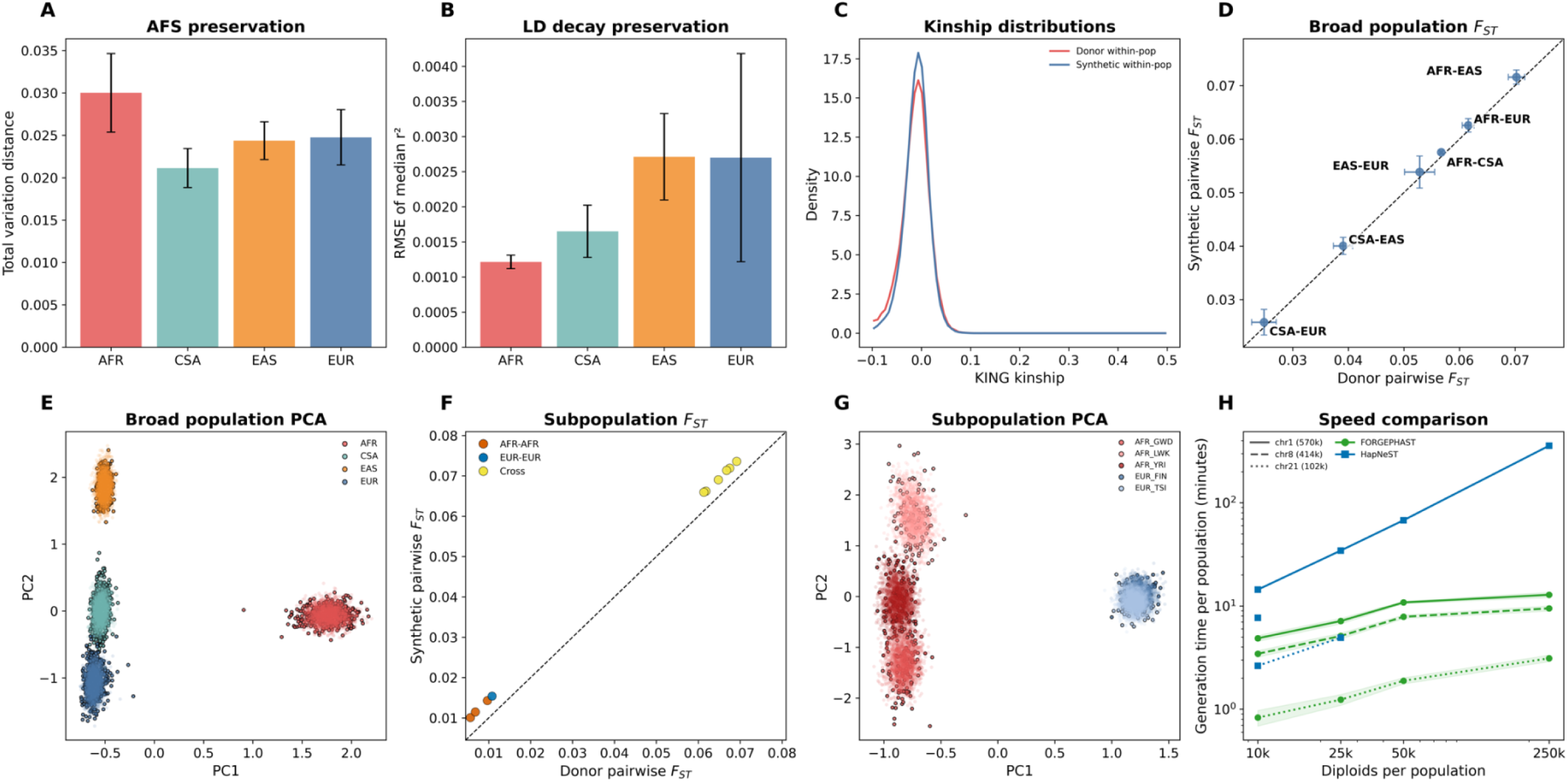
Quality control of homogeneous mosaic synthetic samples against the reference donor panel. Synthetic sample construction and exact benchmark settings are described in the **Supplementary Methods. A:** Total variation distance between donor and synthetic allele-frequency spectra. **B:** LD-decay preservation measured as the RMSE of median *r^2^* between donor and synthetic samples. **C:** KING kinship distributions for donor within-population pairs and synthetic within-population pairs. **D:** Donor-panel pairwise *FS*T versus matched synthetic pairwise *F*ST across broad population pairs. **E:** Synthetic samples, shown as semi-transparent points, projected onto donor-panel PCs and plotted with donor samples, shown as opaque points. **F:** Donor-panel pairwise *F*ST versus matched synthetic pairwise *F*ST for homogeneous EUR and AFR subpopulation simulations; “Cross” denotes between EUR- and AFR-derived subpopulations. **G:** Synthetic subpopulation samples, shown as semi-transparent points, projected onto subpopulation donor PCs and plotted with donor samples, shown as opaque points. **H:** Generation-time comparison (minutes) between HAPNEST and FORGEPHAST. HAPNEST results are shown for chromosome 1, 8, and 21. Chromosomes 8 and 21 are only shown for runs that completed (10K for 8, 10K and 25K for 21); larger runs terminated with non-recoverable segmentation faults under the same 420 GB as the successful chromosome 1 runs.

A compact demonstration of homogeneous subpopulation simulation was accomplished using a donor panel of mixed AFR and EUR subpopulations, consisting of 1KG individuals from GWD (n=113), LWK (n=102), YRI (n=107) African populations and FIN (n=105), and TSI (n=111) European populations. Ten thousand synthetic diploid individuals were created for each population for chromosome 21. 2000 synthetic diploids per subpopulation were subsampled to compute pairwise *F*ST and PCA, of which PC1 and 2 are shown in **Figure 3**, with remaining QC metrics found in **Supplementary Data 1**.

To establish the speed of FORGEPHAST, we iterated across synthetic diploid size 10k, 25k, 50k, and 250k. Comparisons between FORGEPHAST and the program HAPNEST(Wharrie *et al*., 2023) are displayed in **Figure 3H**. FORGEPHAST generated 250, 000 diploids per population for chromosome 1 in under 15 minutes, excluding QC. HAPNEST completed the full size sweep for chromosome 1, the 10, 000-diploid run for chromosome 8, and the 10, 000- and 25, 000-diploid runs for chromosome 21; larger chromosome 8 and 21 runs terminated with non-recoverable segmentation faults. Because chromosome 1, the largest chromosome, completed successfully, these failures were unlikely to reflect insufficient memory allocation. We benchmarked against HAPNEST because it has previously been reported to be faster than other reference-based simulators.

As an example of admixed simulation using the msprime module in FORGEPHAST, we generated populations derived from two- and three- way admixture pulses. Briefly, the two-way admixture was derived from the homogeneous AFR and EUR populations described in **Supplementary Table 2** with a 50/50 ratio. The three-way admixture was simulated using a donor panel consisting of AFR, EUR, and a defined homogeneous AMR panel(Mallick *et al*., 2016) (included groups can be found in **Supplementary Table 2**), at a ratio of 25/65/10. A total of 10k diploids were generated for each for chromosome 21 based on a pulse happening 10 generations previously. Timing for 5k, 10k, 25k, and 100k individuals along with example QC results are shown in **Figure 4**, including empirical tract length vs the expected distribution for both simulations and PC1 and 2 plots for the three-way admixture.

**Figure 4.**
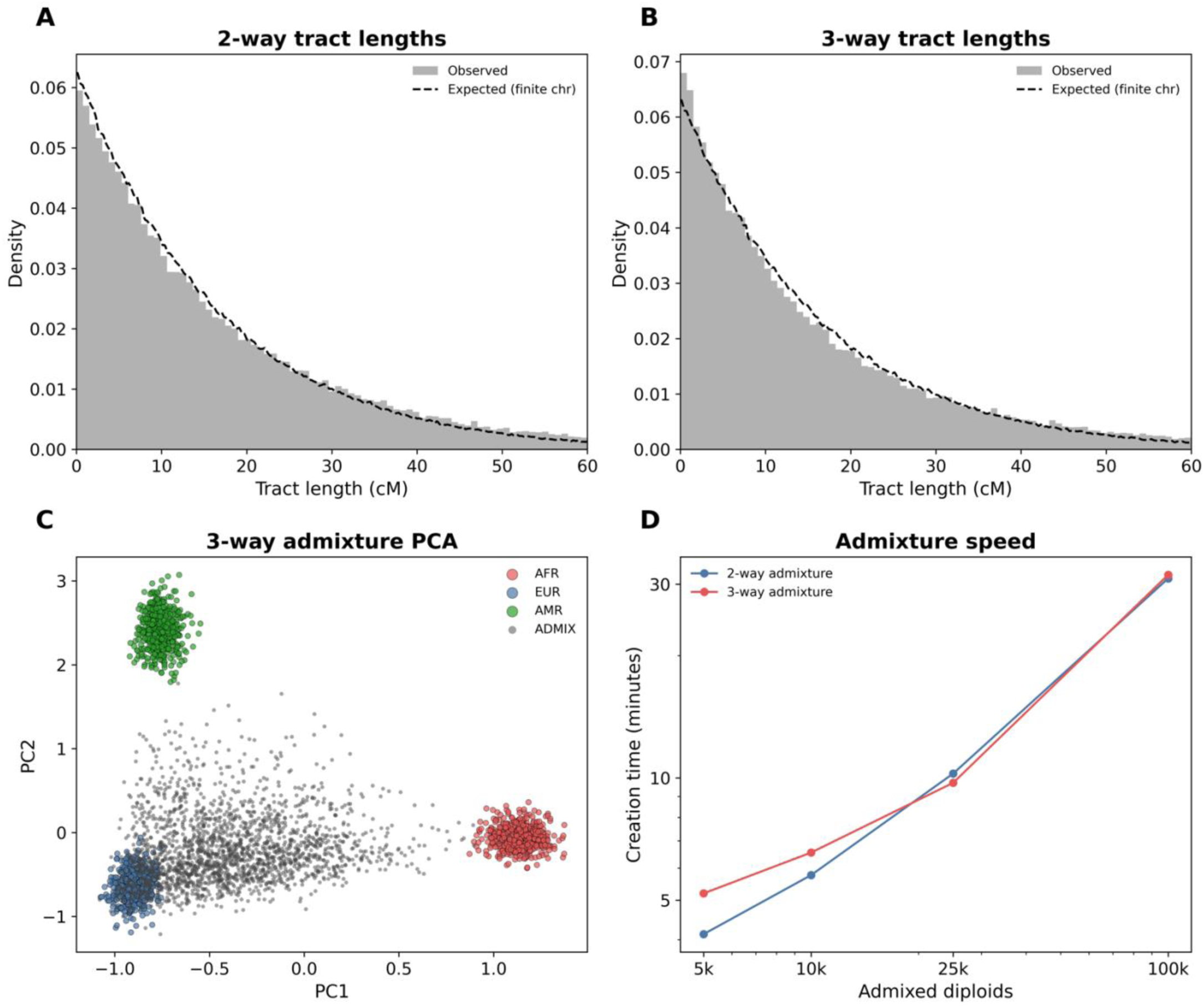
Quality control of two- and three-way admixed synthetic populations. Admixed sample construction, source-population weights, tract-generation parameters, PCA filtering, and benchmark settings are described in the Supplementary Methods. **A:** Observed chromosome 21 local-ancestry tract-length distribution for two-way admixture, compared with the expected finite-chromosome distribution. **B:** Observed chromosome 21 local-ancestry tract-length distribution for three-way admixture, compared with the expected finite-chromosome distribution. **C:** PCA of three-way admixed and non-admixed synthetic comparator cohorts in donor-defined PC space. Principal components were fit on the underlying AFR, EUR, and AMR donor samples. Five hundred non-admixed synthetic comparator diploids per source population and 2,000 three-way admixed diploids were projected into this space; only the projected cohorts are shown. **D:** Creation time for two-way and three-way admixed cohorts across increasing numbers of admixed diploids

### PHENOTYPE SIMULATION

Phenotype simulation in FORGEPHAST is carried out as described above and in the **Supplementary Methods**. To assess calibration, we generated null phenotypes for 150, 000 EUR and 65, 000 AFR synthetic samples, with both fixed and background genetic variance set to zero. Pooled and population-stratified REGENIE analyses using 20 LD-pruned global PCs were well calibrated (λGC = 0.999 pooled, 0.997 EUR, and 0.990 AFR; Figure 5A). REGENIE was used as the primary GWAS engine- REGENIE parameters for this and all subsequent GWAS on synthetic data are described in more detail in the **Supplementary Method**.

**Figure 5.**
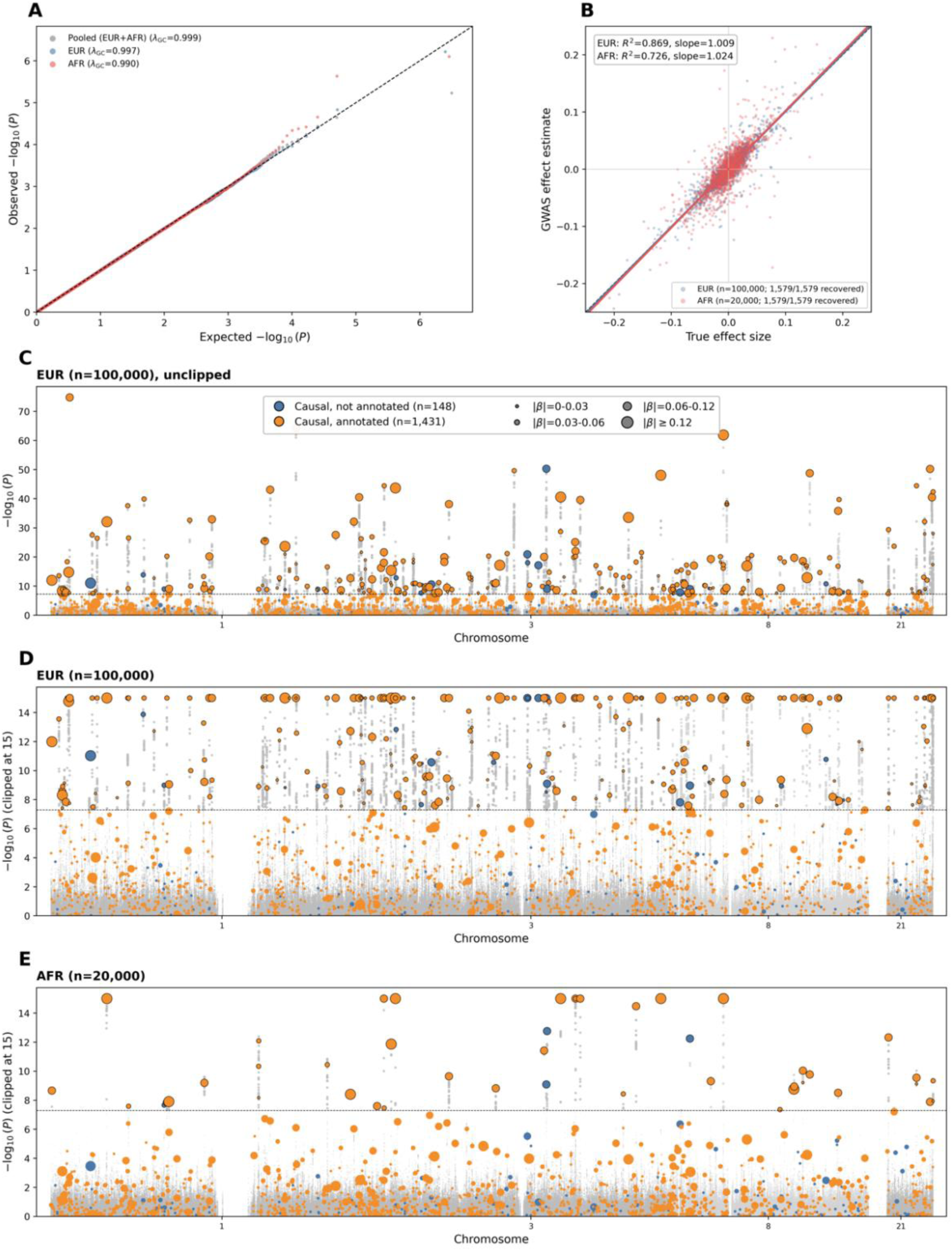
Null phenotype calibration and GWAS of a synthetic quantitative phenotype. **A:** A phenotype constructed with no genotypic effect shows the expected null distribution in the combined EUR/AFR cohort and in population-stratified analyses, as measured by genomic inflation factor. **B:** REGENIE-estimated effect size versus true simulated effect size for causal variants in each population. **C,D:** Manhattan plots for the EUR cohort, shown with an unclipped y-axis in **C** and a clipped y-axis in **D**. Point size denotes the absolute true effect size for causal variants. **E**: Manhattan plot for the AFR cohort, shown with the same y-axis clipping as **D**.

With clear null calibration, we next simulated a phenotype across 100K synthetic EUR diploids and 20K AFR diploids, with detailed description of creation in the **Supplementary Methods**. The phenotype was simulated across 4 chromosomes (1, 3, 8, 21) using an effect size drawn from a Student-t distribution (df=4), with cohort-level *h^2^* =0.25, 0.1% polygenicity, MAF scaling, per-variant population effect size jitter of 0.05 with a cross-ancestry correlation derived automatically for pairwise *F*ST (target correlation 0.94, real correlation 0.95), allowed causal variant MAF ratio of less than or equal to 8, and clump mode with the default gene annotations described above. This resulted in a simulation of phenotypes across 1.58 million candidate variants, after thresholding at a cohort-wide MAF floor of 0.0005; following the MAF ratio filter, this left 800 thousand variants. True per-population statistics, including effect size, phenotypic, and variant distributions are reported in **Supplementary Table 3**.

Population-stratified GWAS was performed with REGENIE on the simulated phenotype. Under the simulated genetic architecture (cohort-level *h*² = 0.25), the EUR cohort exhibited λGC = 1.40 and the AFR cohort exhibited λGC = 1.09 (**Supplementary Figure 2**), consistent with the large disparity in cohort size (n = 100, 000 vs. n = 20, 000). **Figure 5B** compares REGENIE-estimated and true simulated effect sizes for both populations. **Figures 5C-D** show EUR Manhattan plots with unclipped and clipped y-axes, respectively, and **Figure 5E** shows the AFR Manhattan plot with the same clipped y-axis scale as **Figure 5D**, highlighting the decreased power in the AFR cohort.

### POLYGENIC RISK SCORE DEMONSTRATION

As a demonstration of a use case for FORGEPHAST, we applied seven modern PRS methods to a split AFR/EUR biobank-simulation of chromosome 1, 3, 8, and 21 using the cohort and variants described above. We specifically chose PRS methods that were created with the intention of cross-population ancestry transfer: PRS-CSx(Ruan *et al*., 2022), JointPRS-auto(Xu *et al*., 2025), CT-SLEB(Zhang *et al*., 2023), PROSPER(Zhang *et al*., 2024), SDPRX(Zhou *et al*., 2023), X-Wing(Miao *et al*., 2023), and MUSSEL(Jin *et al*., 2024). We also included population-stratified C+T as a baseline comparator(Privé *et al*., 2019).

Our simulated biobank was generated using FORGEPHAST’s mosaic model simulation as described in detail in the **Supplementary Methods** and was divided into 120K individuals in train, and 50K each in validation and test cohorts for a total of 100K individuals in the evaluation cohort. Train consisted of 100K synthetic EUR diploids and 20K AFR diploids, while both validation and test were evenly divided between the two populations.

FORGEPHAST was used to generate continuous phenotypes across a grid of different simulations parameters. The parameters varied were *h*^2^ (selected values: 0.05, 0.25, 0.5), polygenicity (0.0005, 0.001, 0.005), allowed MAF ratio differences between EUR and AFR (2 [“tight”], 8 [“moderate”], 300 [“extreme”]), and whether effect sizes could vary or not between populations (jittered/unjittered). Because h^2^ is nominally targeted on a cohort-level instead of population-level, realized h^2^ can differ between populations, particularly when MAF varies greatly between populations for a causal variant, as in the extreme regime. These realized h^2^ values are shown in **Figure 6A** for the 54 simulated phenotypes. Detailed information about both genotype and phenotype simulation are deferred to **Supplementary Methods**.

**Figure 6.**
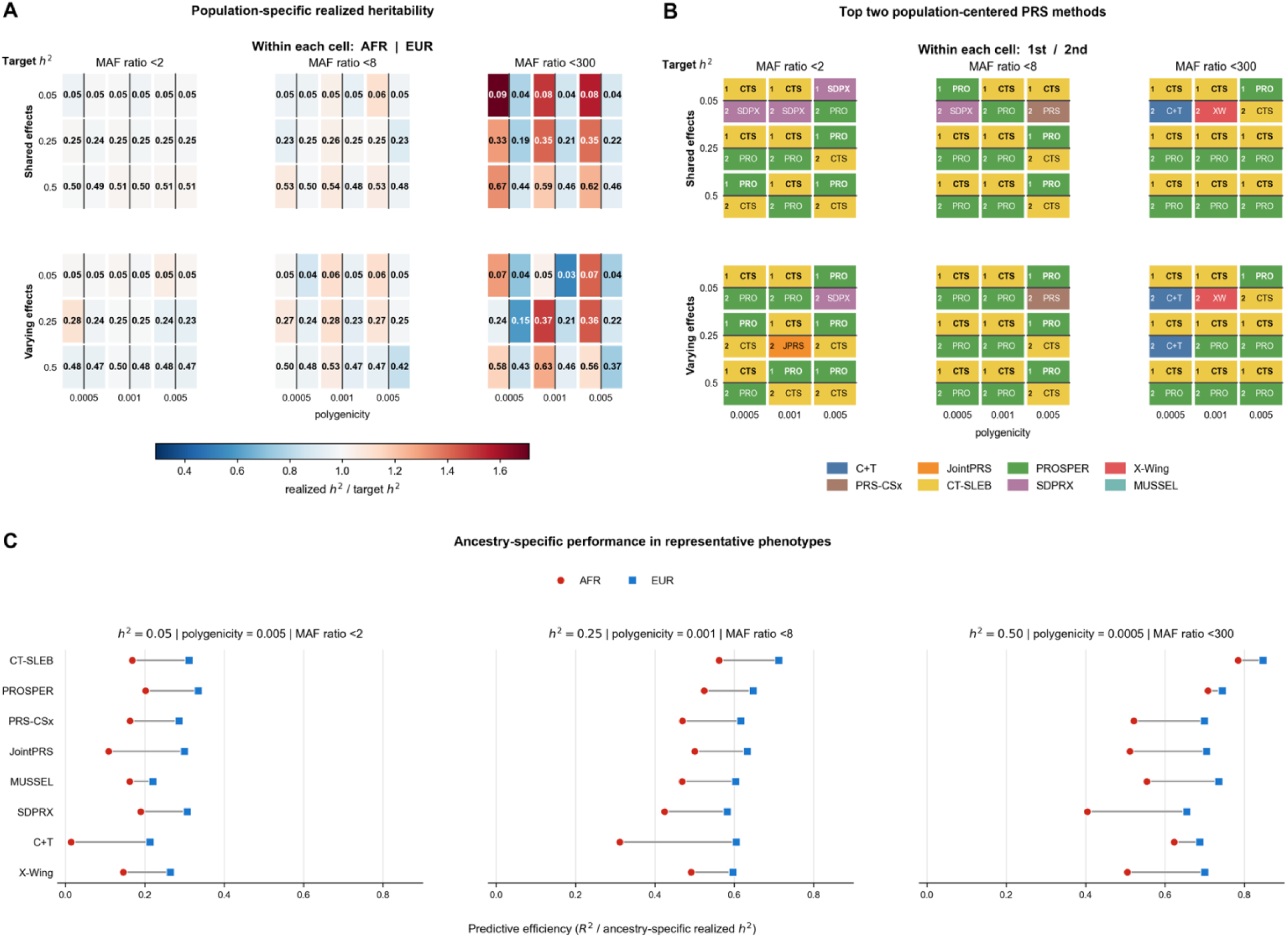
Polygenic risk score tool demonstration across 54 phenotype grid of split EUR/AFR cohort. **A:** Population-specific realized heritability across all 54 phenotypes. Cell color represents realized *h*² divided by the target cohort *h*², and labels report realized *h*². **B:** Top two PRS methods by population-centered test R² for each phenotype. The top half of each cell shows the best-performing method and the bottom half shows the second-best method. **C**: For 3 representative phenotypes, ancestry-specific test R² divided by ancestry-specific realized *h*² is shown for AFR and EUR for all 8 PRS methods.

GWAS summary statistics were generated using REGENIE in a population stratified-method, with the null model utilizing 100 PCs as covariates. These were generated per-population. Detailed results of each GWAS are presented in the **Supplementary Data 2**. Each PRS method was run in its standard mode as described in respective publications or software package, with a grid search over parameters if necessary. Additional details about the GWAS construction and PRS parameters and methodology is supplied in the **Supplementary Methods**. Of note, PRS-CSx, SDPRX, X-Wing, and Joint-PRS are hard-coded to only accept or were not originally conceived with the ability to extend beyond HapMap3 variants (Xu *et al*., 2025). We retained these restrictions rather than extending the methods beyond their documented variant universes. Consequently, this analysis compares complete method implementations, including their supported variant sets, rather than statistical models evaluated over an identical set of variants.

Results of the 54 phenotypes are displayed as top 2 winners by population-centered R^2^ in the test cohort in **Figure 6B**. Group-centering removes this intercept-like component and evaluates whether a PRS captures within-population genetic variation, which is the relevant transferability question in this benchmark. Full results across all methods for the top-validated parameters are presented in the **Supplementary Data 2.**

Performance across the 54 architectures was concentrated among CT-SLEB and PROSPER (Figure 6B). CT-SLEB produced the highest population-centered R^2^ in 37 phenotypes, PROSPER in 16, and SDPRX in one. CT-SLEB and PROSPER also occupied 95 of the 108 top-two positions. These results reflect both model construction and variant coverage. CT-SLEB and PROSPER can construct scores from broader ancestry-aware variant sets, whereas several competing workflows are restricted to HapMap3 variants.

PROSPER was most competitive under highly polygenic architectures with relatively similar causal-variant frequencies. This pattern is consistent with PROSPER’s combination of population-specific penalized regression and cross-population shrinkage: when causal variants are represented at comparable frequencies in both populations, information can be shared across GWAS with less risk of shrinking toward poorly supported effects. In contrast, CT-SLEB dominated the extreme MAF-ratio conditions, winning 16 of 18, consistent with its ancestry-specific variant selection and effect estimation retaining signals that differ in informativeness across populations.

Population-specific evaluation recapitulated the expected cross-ancestry transferability gap (example phenotypes in **Figure 6C**; full grid in **Supplementary Figure 3A**). Neither CT-SLEB nor PROSPER showed a disproportionate cross-ancestry transferability penalty. Across all 54 architectures, their median normalized AFR-to-EUR performance declines were similar (22.4% and 22.6%, respectively) and smaller than those observed for most other methods. The magnitude of the transferability gap was instead strongly architecture-dependent, increasing under lower heritability, higher polygenicity, and greater cross-population allele-frequency divergence. In 17 of 54 phenotypes, the top-performing EUR and AFR methods were different. CT-SLEB was the ancestry-specific winner in 41 EUR and 29 AFR evaluations, whereas PROSPER won 10 EUR and 23 AFR evaluations. PROSPER’s overperformance in AFR was mostly concentrated in smaller MAF ratios, highlighting its capacity to share information between ancestries again (**Supplementary Figure 3B**).

## DISCUSSION

Here, we present FORGEPHAST, a simulation toolkit for generating diverse genotype and complex phenotypes, as well as a companion storage format termed HapStore. While other tools may have individual parts of FORGEPHAST, as well as potential additional components, FORGEPHAST combines all of its features together in an end-to-end runnable pipeline (**Table 1**). We provide a fully integrated QC suite for both the genotype and phenotype portions of the toolkit, without requiring significant intervention to derive important measures. This toolkit is written completely in Python and is easily extendible. YAML configuration files and CLI modules provide easy plug-and-play access to the toolkit overall.

**Table 1.** Comparison of FORGEPHAST with genotype and genotype-phenotype simulation frameworks. A filled cell denotes that the feature is described in the original publication, source code, user-facing documentation, or a subsequent demonstrated use. An “X*” denotes that the capability is available through package-native components or minimal wrapper code, but is not exposed as a single first-class workflow. Blank cells indicate that direct evidence for that feature was not identified. Demonstrated biobank-scale simulation is defined as a documented simulation of at least 100,000 diploid individuals

| Method/Tool<br>(Citation) | Reference-panel<br>haplotype<br>amplification | User-defined<br>population<br>panels | Admixed<br>local-ancestry<br>tracts | Demographic<br>and population<br>parametrized<br>model | Integrated<br>genomic<br>QC | Integrated<br>phenotype<br>simulation | Local-ancestry<br>specific<br>effects | Phenotypic<br>evaluation and<br>QC | Haplotype -<br>resolved<br>synthetic<br>storage | Matrix-<br>native<br>analysis<br>storage | Demonstrated<br>biobank-scale<br>simulation |
| --- | --- | --- | --- | --- | --- | --- | --- | --- | --- | --- | --- |
| FORGEPHAST | X | X | X |  | X | X | X | X | X | X | X |
| HAPNEST (Wharrie<br>*et al.*, 2023) | X | <sup>a</sup> |  | X<sup>b</sup> | X | X |  | X |  |  | X |
| HAPGEN2 (Su *et al.*,<br>2011) | X | X |  |  |  | X |  |  | X |  | X<sup>c</sup> |
| tskit ecosystem (see<br>below<sup>d</sup>) |  | X* | X* | X | X* | X | X* |  | X | X* | X |
| Admix-kit (Hou *et al.*,<br>2024) | X* | X | X | X | X* | X | X |  | X | X* |  |
| G2P (Tang and Liu,<br>2019) | <sup>e</sup> | X |  | X* | X | X |  | X |  |  | X |
| HAP-SAMPLE2<br>(Sun *et al.*, 2025) | X | X |  | X<sup>f</sup> |  | X |  |  | X* |  |  |

<sup>a</sup>HAPNEST supports custom population structures by assigning segment weights among fixed superpopulation labels, but the implementation is constrained to the built-in 1KG+HGDP superpopulation set; we therefore do not count this as arbitrary user-defined reference panels. <sup>b</sup> HAPNEST includes mutation-age and population-specific parameters, so we count it as limited demographic support rather than arbitrary user-defined demographic-history simulation. <sup>c</sup>HAPGEN2 was included in the HAPNEST benchmark at 100,000 synthetic samples and has also been used for 100,000-individual HAPGEN2 chromosome 1 simulated datasets in FlashPCA2(Abraham *et al.*, 2017). <sup>d</sup>“tskit ecosystem”(Kelleher *et al.*, 2019) refers to the combined tree-sequence ecosystem, including msprime(Baumdicker *et al.*, 2022), SLiM(Messer, 2013)/stdpopsim(Gower *et al.*, 2025), tskit storage/statistics(Ralph *et al.*, 2020), and tstrait-style phenotype simulation(Tagami *et al.*, 2024). <sup>e</sup>G2P includes genotype simulation by sampling genomic blocks from ancestor genotypes, but this is genotype/block-level simulation rather than phased reference-panel haplotype amplification. <sup>f</sup>HAP-SAMPLE2 models specified admixture proportions and recombination from phased haplotypes; we count this as an admixture/resampling generative model.

FORGEPHAST uses well-validated methods to produce haplotypes. Simple demographic models are used; while this may not fully model modern populations, keeping models with fewer hyperparameters allows for a potentially larger selection of populations without having to define population-specific effective population sizes and recombination rates, for instance. While FORGEPHAST can maintain most markers of genetic fidelity between donors and synthetic samples, Hardy-Weinberg equilibrium emerges in greater agreement for synthetic samples. Similarly, other markers of genetic similarity show slight reductions in synthetic samples. This is to be expected given that Li-Stephens and our donor mosaic model closely approximates Hardy-Weinberg assumptions, without any type of selective pressure and the cryptic relatedness found in real populations.

Several limitations of the synthetic genotype framework should be noted. The admixture model does not capture the more complex demographics arising from continuous gene flow, multiple admixture waves, or sex-biased migration. Extending the demographic model to support these scenarios is a natural direction for future work. Similarly, the Li–Stephens mosaic copying engine is neutral by construction: it preserves linkage disequilibrium patterns from the donor panel but does not model natural selection. Simulated LD around causal loci therefore lacks signatures of selection, which may be relevant for fine-mapping benchmarks though less so for PRS evaluation. Additionally, while common-variant LD fidelity is well preserved, the finite donor pool limits the ability to generate realistic rare-variant site frequency spectra. This is a limitation of FORGEPHAST; FORGEPHAST primarily targets common-variant polygenic association and scoring applications rather than de novo or ultra-rare variant architectures, for which demographic generative simulators may be more appropriate.

The PRS demonstration showed that apparent method performance depends on phenotype architecture, population representation, and the variant universe available to each workflow. Although this experiment varied phenotype architecture rather than demographic history, FORGEPHAST can vary both axes. The simulations recapitulated the well-described cross-ancestry transferability gap (Kachuri *et al*., 2024) and provide a controlled framework for distinguishing contributions from GWAS sample size, variant ascertainment, allele-frequency and LD differences, model conditioning, and assumptions about cross-population effect sharing. Because the present comparison used each method’s supported variant universe, it evaluates complete implementations rather than statistical models under perfectly matched inputs, which is a known real-world limitation of some of these methods (Xu *et al*., 2025). Real traits may additionally involve dominance, epistasis, or gene-environment interactions not modeled in this initial release. However, many of these effects have been shown to contribute to variance at lower levels and were not modeled in this initial release(Hill *et al*., 2008).

HapStore addresses an emerging infrastructure gap as biobank-scale analysis moves to cloud environments. Its competitive performance against PLINK2’s compiled C++ kernels for operations such as PCA, LD correlation, and KING kinship reflects the advantage of operating directly on the analysis-native haplotype representation without format conversion, rather than a general language-level performance claim. As cloud-native genomics platforms mature, storage formats that support direct programmatic access from scientific computing and machine learning frameworks will become increasingly important for method development workflows.

FORGEPHAST integrates capabilities that currently require assembling multiple separate tools, including reference-based haplotype synthesis, admixture tract simulation, local-ancestry-aware phenotype generation, and population-genetic QC, into a single configurable pipeline, with HapStore providing a storage layer suited to cloud-native and ML-oriented analysis workflows. To lower the barrier to diverse simulation for the community, we release pre-generated synthetic datasets of homogeneous and admixed populations with paired ground-truth phenotypes in HapStore format. As PRS methods increasingly target admixed and underrepresented populations, benchmarking platforms that provide controlled ground truth across the full chain will be essential for distinguishing genuine methodological advances from artifacts of evaluation design.

## Acknowledgements

Generative AI tools, including OpenAI Codex and Anthropic Claude, were used to assist with code development, documentation, consistency checking, figure preparation, and language editing. All resulting code, analyses, figures, and text were reviewed and validated by the authors, who take full responsibility for the work.

WC: ~4700

## Supplementary Methods

Scripts and source data required generally to generate data, comparisons, and figures can be found in this paper’s Github at https://github.com/sriraj-lab/forgephast-manuscript-figures.

The following sections describes modeling details used in FORGEPHAST. Technical documentation may be found in FORGEPHAST’s git repository.

### Mosaic Donor Model

The mosaic donor model uses an adaptation of the Li-Stephens model(Li and Stephens, 2003) to assign switch probabilities and then draws a new haplotype from the donor pool. The following mathematical model is used; in this model, D is the genomic distance in centimorgans from a marker *j* and *τ* is the mean tract length:

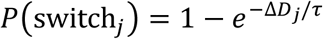

Probabilities of switching to another haplotype are constant across all donor haplotypes in unweighted mode. However, in the weighted mosaic donor model, the weight of each source population, *Wk* is used to alter the selection by splitting the specified weight across all individuals of that source population. This can be used to enrich or deplete a certain source population compared to the donor pool. In both modes, if the sampled donor is the same as the current donor, the next-indexed valid donor is selected instead. All simulations used the Beagle-hosted GRCh38 maps, found at https://faculty.washington.edu/browning/beagle/beagle.html.

### Mosaic Donor QC

FORGEPHAST implements many QC metrics to validate that synthetic populations genetically match their donor populations. Each method uses custom Python kernels that interface closely with HapStore as described here. Default parameters for the gates are given in **Supplementary Table 1**.

#### FST

We utilize Hudson’s *F*ST^(^Hudson *et al*., 1992^)^ to determine if populations separate similarly in donor and synthetic samples, as well if random subsamples of synthetic samples mimic the donor-synthetic sample similarity. We reimplement the PLINK2-style Hudson *F*ST estimation(Chang *et al*., 2015): for a given variant, *j*, with alternate allele frequencies in populations 1 and 2 equal to *p1, j* and *p2, j* and total chromosome count *n1, j* and *n2, j*, we compute:

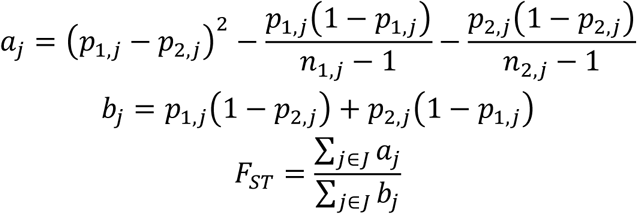

FORGEPHAST’s initial gate compares between-population pairwise Hudson *F*ST among synthetic populations to the corresponding donor-panel pairwise *F*ST for the same population labels. For each population pair, the relative change is defined as |*F*ST(synthetic) - *F*ST(donor)| / |*F*ST(donor)| when the donor baseline is nonzero; when the donor baseline is effectively zero, the absolute difference |*F*ST(synthetic) - *F*ST(donor)| is used instead. As an internal homogeneity check, each synthetic population was also split into two subsets and *F*ST was computed between subsets; runs failed this check if the absolute within-population split *F*ST exceeded the configured threshold.

Because synthetic cohorts may be generated at scales larger than the finite donor panels from which they are copied, FORGEPHAST additionally evaluates pairwise *F*ST deviations against a donor-resampling null. For each population pair, the observed statistic is the absolute difference between synthetic pairwise *F*ST and the corresponding full-donor pairwise *F*ST. FORGEPHAST then repeatedly resamples donor individuals at the smaller of the synthetic QC sample size or donor sample size, recomputes pairwise *F*ST, and estimates the fraction of donor-resampling replicates with a mean absolute pairwise *F*ST deviation at least as large as the observed synthetic deviation in eligible population pairs. The stochastic *F*ST gate passes when this empirical probability is greater than or equal to the configured minimum value.

#### KING

As an orthogonal approach to *F*ST, the KING-robust kinship estimator was used following Manichaikul et al’s formulation(Manichaikul *et al*., 2010):

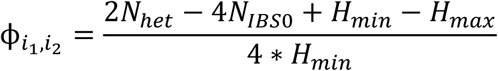

This is an individual pairwise estimator. *N*_ℎ*et*_ is the number of loci where both individuals are heterozygous and *N_IBS_*_0_ are the loci which are opposite homozygous. Each of the *H* terms are the minimum/maximum heterozygote counts across the two samples.

FORGEPHAST uses KING to summarize pairwise relatedness structure within and between simulated populations. For each population, KING-robust kinship is computed for sampled pairs of donor individuals and sampled pairs of synthetic individuals, yielding within-donor and within-synthetic kinship distributions. These distributions are compared by summary statistics and overlaid histograms to assess whether amplification preserves within-population relatedness structure. Separately, FORGEPHAST computes KING-robust kinship for sampled pairs of synthetic individuals drawn from different synthetic populations. The pooled within-synthetic and cross-synthetic distributions are compared by ROC AUC, providing a kinship-based diagnostic of population separation in the synthetic cohort.

#### LD decay

FORGEPHAST computes LD using the *r*^2^ formulation over variant pairs within a centimorgan window for a sampled donor and synthetic set of haplotypes, filtering for minimum MAF. We determine weighted RMSE of median *r*^2^ between synthetic and donor. As additional shape diagnostics, FORGEPHAST computes the AUC of the median *r*^2^ and the half-decay distance, which is defined as the smallest genetic-distance bin midpoint at which median *r*^2^ falls below a fixed threshold. The LD gate passes when the weighted RMSE is below the configured threshold and the donor/synthetic relative half-decay-distance difference is below the configured threshold; if both curves never cross the half-decay threshold, the half-decay component is treated as passing. AUC was reported descriptively.

#### PCA

PCA was used as a primary visual check that synthetic cohorts preserved donor-panel population structure. For each chromosome-level QC run, FORGEPHAST first constructed a shared donor/synthetic variant axis by intersecting donor and synthetic variants by position, then sampled up to the configured maximum number of variants and diploid individuals per population. PCA variants were filtered using donor allele frequencies, requiring finite allele frequency, chromosome count at least four, complete donor and synthetic genotypes, nonzero variance, and minor allele frequency at least the configured threshold. Genotypes were mean-centered and variance-scaled using donor allele frequencies. Principal components were fit on the donor genotypes only, and synthetic genotypes were projected into this donor-defined PC space. The first two projected PCs were used for visual inspection. For quantitative comparison, FORGEPHAST computed pairwise population-separation AUCs in donor space and synthetic-projected space, using the donor population-centroid contrast as the scoring direction. The absolute donor/synthetic AUC difference was reported for each population pair.

#### Allele Frequency Spectrum

Allele frequency spectrum (AFS) QC compared donor and synthetic MAF distributions within each population. FORGEPHAST constructs histograms of these distributions and reports total variation distance, Jensen-Shannon divergence, and KS distance as diagnostics.

#### Secondary metrics

As secondary diagnostics, FORGEPHAST also reports runs of homozygosity (ROH), Hardy-Weinberg equilibrium (HWE) statistics, and the inbreeding coefficient *F* which are commonly used in cohort-level genotype QC. ROH was summarized per individual as total ROH length, number of segments, and maximum segment length using configurable minimum segment length and maximum inter-marker gap thresholds. HWE was evaluated from per-variant genotype counts after minimum MAF and sample count filters. Inbreeding *F* was computed from genotype-count summaries at filtered variants and compared between donor and synthetic populations using distributional summaries. These metrics were treated as qualitative diagnostics by default rather than primary pass/fail gates, because the mosaic generator is not intended to reproduce all ascertainment, relatedness, and cohort-history features that can affect these quantities in real donor panels.

### Admixed Donor Model

Admixed synthetic populations are generated using FORGEPHAST’s msprime-backed msprime_tracts engine. The ancestry process follows the DTWF model implemented in msprime(Baumdicker *et al*., 2022). Users specify source populations, admixture proportions, the time since the pulse-admixture event, and the number of synthetic individuals. In FORGEPHAST, for each admixed population, users specify one or more source populations and source weights that sum to one. For multi-source populations, FORGEPHAST simulates a pulse-admixture ancestry process in msprime using the user-supplied genetic map and admixture time in generations. The msprime simulation is used to generate local-ancestry tract boundaries and source-population labels, rather than sequence variation from the donor individuals themselves. User-specified inputs include the source populations and admixture proportions, the pulse-admixture time, and the number of synthetic individuals. The current implementation uses msprime’s DTWF ancestry model to generate local-ancestry tract structure, with internal source and target population sizes fixed at 10, 000. Each ancestry-labeled tract is then instantiated by sampling a donor haplotype segment from the corresponding aligned source panel. Thus, admixed genomes contain observed donor haplotype segments arranged according to msprime-generated ancestry tracts, without simulating new mutations. Continuous migration and other more complex demographic models are not currently represented in this admixed-generation mode.

### Admixed Donor QC

For admixed cohort QC, FORGEPHAST relies primarily on deviation from expected mixture under the model parameters. First, FORGEPHAST compared genome-wide observed source proportions to the configured admixture weights. Second, local-ancestry tract lengths were compared to the finite-chromosome expectation implied by the pulse-admixture time and chromosome genetic length(Liang and Nielsen, 2014). Crucially, this deviates from the expected plain infinite-chromosome exponential with mean tract length 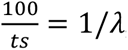, where *t* is the time since pulse and 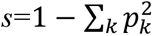 is the probability that adjacent ancestries differ given by the mixture weight. Instead, under a finite-chromosome Poisson-switch distribution, the CDF and expected tract length for chromosome of length *L* in centimorgans is equal to:

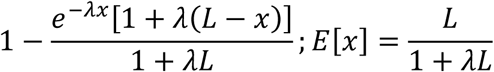

This check evaluates whether recombination has produced the expected tract scale rather than only the correct genome-wide ancestry fraction. Third, FORGEPHAST compared the variance of per-individual ancestry proportions to the corresponding pulse-admixture expectation, because a cohort can match the target mean ancestry while having incorrect individual-level ancestry dispersion. Finally, admixed synthetic samples were projected into donor-fit PCA space alongside the source populations and inspected for the expected interpolation between source panels.

### Phenotype Generation

FORGEPHAST’s phenotypic generation is very user-parametrizable to model a vast number of complex phenotypes under various scenarios. Specific implementation details can be found in the code; here, we provide a compact summary of some crucial details.

#### Inputs and cohorts

FORGEPHAST’s phenotype generation is HapStore native and uses HapStore diploid genotypes, cohort sample list, population labels, and population tracts if local ancestry-based effect sizes are configured. Population labels do not solely need to be ancestral labels; any division of a cohort can be used to accentuate phenotypic modeling as required.

#### Causal variant selection

FORGEPHAST can rapidly define the causal-variant universe by scanning HapStore genotype blocks, computing pooled and source-population MAFs, and retaining variants above a configured pooled MAF threshold. This table contains variant identifiers and population-specific allele-frequency annotations. FORGEPHAST first filters this pool to the configured MAF range and assigns variants to user-defined MAF buckets. Within each bucket, the simulation can include both a fixed causal set, shared across phenotype replicates, and a replicate-specific subset sampled without replacement.

Sampling probabilities can optionally be weighted by annotations. In the default gene-window mode, variants falling within annotated gene intervals (GENCODE v49(Frankish *et al*., 2023)), including the configured upstream region, receive an increased sampling weight; custom BED annotations can also be supplied. To avoid selecting variants with extreme cross-population allele-frequency shifts unless explicitly desired, FORGEPHAST also supports a “tightness” filter based on the maximum pairwise MAF fold-change across configured populations.

#### Effect size model

After variants are selected, effect sizes can be generated from different distributions: normal, student’s T, Laplace, and a mixture of normals. To reflect the general understanding that rarer variants have larger functional effects(Schoech *et al*., 2019), MAF scaling can be applied as functions of 2*p*(1-*p)*. Effect size scaling is based on an “anchor” MAF-this defaults to an anchor population, but these can also be defined for the full cohort or custom populations as needed. Furthermore, effect sizes are allowed to vary between ancestries based on a difference from a shared base. The overall correlation of these effects can be manually specified or inferred from an exponential decay transform of *F*ST.

#### Local-ancestry effects

Hou and colleagues modeled allelic effects according to the local-ancestry background on which each allele occurs and found that these effects were highly correlated across ancestry backgrounds (Hou *et al*., 2023). FORGEPHAST supports this model by using local-ancestry tracts to assign the corresponding ancestry-specific causal effect to each allele.The genetic component is then summed over haplotypes and causal variants for an individual. Thus, an individual who has two copies of the causal allele from different reference backgrounds could potentially have two separate effects summed together, if ancestral effect size variability is enabled.

#### Phenotype variance budget

The central control of heritability (*h^2^*) is controlled by a phenotype variance budget, where the phenotype is composed of additive components:

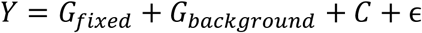

The fixed genetic component is realized from the effect sizes of causal variants; the background effect can either be set to none, GRM-derived, or from pre-computed kinship and is included to mimic infinitesimal architectures. To reach the necessary *h^2^*, FORGEPHAST computes the raw fixed genetic component and rescales it with a single cohort-level multiplier to match the configured fixed-effect heritability target. The same multiplier is applied to the stored effect-size truth table, so reported effect sizes correspond to the generated phenotype. Covariate and residual effects are also scaled to properly reflect their contribution to the variance budget.

### Binary traits

For binary traits, FORGEPHAST first generates the same continuous value used for quantitative phenotypes and uses that as a liability. To allow prevalence to be standardized, FORGEPHAST standardizes (mean-centered, standard deviation of 1) the liability within the simulated cohort, such that each individual has a standardized liability, *zi*. Disease probability is then assigned using a logistic liability model(Falconer, 1965)

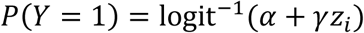

For a single target prevalence, the intercept, *α*, is numerically calibrated so that the mean predicted probability equals the requested prevalence. When population-specific prevalences are specified, FORGEPHAST solves separate source-population intercepts *α_a_* so that each source population matches its target prevalence in expectation.

For admixed individuals, the intercept is not separately calibrated to a fixed admixed prevalence; instead, source-population intercepts are combined according to the individual’s local-ancestry exposure in the causal architecture, and disease status is sampled from the resulting probability. That is, each admixed individual has a separate intercept determined by the fraction of local-ancestry exposure. Thus, admixed prevalence arises from the specified source-population prevalences, local ancestry, effect architecture, and liability distribution rather than being imposed directly.

FORGEPHAST preserves the full simulation truth, including causal-variant membership, population-specific effect sizes, realized component variances, and summary measures of cross-population effect geometry. These outputs allow downstream GWAS and PRS results to be compared directly against the known generative architecture.

### HapStore Construction

HapStore stores phased diploid haplotypes in a Zarr-backed array format designed for random access along both the sample and variant axes. In this study, HapStore datasets were constructed from statistically phased VCFs or PLINK2/PGEN inputs, with one chromosome-level store created per input chromosome before downstream merges or exports. Each store contains a dense haplotype array encoded as unsigned 8-bit integers, with two haplotypes per diploid sample, alongside separate sample and variant metadata tables. Missing alleles are encoded as 255. Sample metadata preserve sample identifiers, while variant metadata preserve chromosome, position, variant identifier, reference allele, alternate allele, and related annotation fields needed for downstream alignment. Haplotypes are encoded as allele dosages on the haplotype axis, with allele metadata retained so that exports to VCF or PGEN can preserve variant identity and allele orientation. For benchmark datasets, arrays were written with fixed sample and variant chunks and Zstd compression; chunk sizes were chosen to balance sequential scan performance with random sample/variant slicing. Chromosome-level HapStores can be concatenated along the variant axis for genome-wide analyses, while independently simulated or ingested cohorts can be merged along the sample axis when their variant axes are aligned. Monomorphic variants are retained in HapStore unless explicitly filtered for downstream tools.

### HapStore/Python vs PLINK2 Parity and Benchmarking

Timed benchmarks were generated using a synthetic chromosome 5, containing 600k diploid genomes spanning EUR, AFR, CSA, and EAS populations(1000 Genomes Project Consortium *et al*., 2015), with 482k variants, subsetted to the numbers reported in **Figure 2**. For all benchmarks in **Figure 2**, each tool was given a total budget of 16 CPUs. For all figures in **Figure 2**, each method was given a single warm-up run and then a total of 5 independent runs were used as the main timing surface; mean timings and standard error are displayed for each timing. For PCA, LD correlation, and KING, minimum MAF was set at 0.05, with the max number of variants set as 20k, 10k, and 20k, respectively, for the larger sized populations. For KING, up to 1, 000 within-population pairs per population and 1, 000 cross-population pairs per population pair were sampled at 10, 000 individuals; these limits increased to 5, 000 pairs at 50, 000 individuals. KING was not run for the 100, 000-individual condition.

Parity values in Supplementary Figure 1 were computed on chromosome 21 using matched PLINK2 functions and HapStore-backed FORGEPHAST Python kernels. Comparisons used 5, 000 sampled positions, 200 donor and 200 synthetic individuals per population for AFS, and 1, 000 sampled pairs per within-population and cross-population comparison for KING. AFS histogram L1 distances were zero for every donor and synthetic population comparison. KING mean and maximum absolute kinship differences were all below 10⁻⁶.

### Synthetic Population Donor Panel

For figure 3 and 4, donor haplotypes were derived from a phased 1000 Genomes v3 data, grouped into broad, non-admixed source panels. The composition of each population and what subpopulations were included is reported in **Supplementary Table 2.** For the homogeneous synthetic populations, AFR, CSA, EAS, and EUR donor panels were used, while the admixed synthetic populations used AFR, AMR, and EUR donor panels.

### Homogeneous Synthetic Population Generation

Synthetic populations were generated using the unweighted donor model for chromosome 1, 8, and 21. For the QC panels, 10, 000 diploid individuals (20k haplotypes) were generated per source panel. Mean tract length was set at 1.0 cM. Extended QC metrics are presented in **Supplementary Data 1**. The timing figure was carried out in a similar fashion to the other generation. Benchmarks were run on a single high-memory HPC node using 48 CPUs and 420 GB allocated memory. This allocation was used as a standardized benchmark envelope across chromosome and sample-size conditions; timings report generation plus merge time and exclude QC. Panels A, B, and D show means across chromosomes 1, 8, and 21, with error bars denoting standard deviation across chromosomes.

Subpopulation preservation was evaluated using a five-way chromosome 21 synthetic cohort generated with the FORGEPHAST mosaic donor model to generate 10, 000 diploid synthetic individuals per subpopulation. Donors were restricted to five 1000 Genomes subpopulations: GWD, YRI, and LWK from AFR, and TSI and FIN from EUR, as described in **Supplementary Table 2**.

HapNeST(Wharrie *et al*., 2023) benchmarks were run using the Singularity image *intervene-synthetic-data_latest.sif*, built from the Docker image *docker://sophiewharrie/intervene-synthetic-data*. The same chromosome backbones were used but were retargeted to the HapNest preprocessed input format. We followed the HapNeST GitHub/README pipeline and configuration structure, using the shared preprocessed HapNeST inputs, and modified only the target chromosome, synthetic sample size, and requested source populations. HapNeST was run on equivalent resources to FORGEPHAST: 48 CPUs and 420 GB memory. For HapNeST, the four-population job runtime was divided by four to give generation time per population. In panel H, FORGEPHAST lines and ribbons show the mean ± standard deviation across four source populations; the HapNeST chromosome 1 line represents a single four-population run normalized to time per population.

### Admixed Population Generation

Admixed populations were generated using the msprime-backed engine from 3 donor pools (EUR, AFR, AMR), as mentioned in **Supplementary Table 2**. A total of 10, 000 diploid individuals for chr21 were simulated for 10 generations, following a single pulse. Each ancestry-labeled tract was copied from the corresponding source donor panel. 500 non-admixed synthetic diploids were generated as comparator/QC cohorts and were not used as the donor pool for admixed generation. For 2-way admixture in **Figure 4**, the weights were set evenly for EUR and AFR. For 3-way admixture in **Figure 4**, weights were set to 65/25/10 for EUR, AFR, and AMR, respectively.

For Figure **4C**, principal components were fit on 1, 180 source-panel donors (512 AFR, 522 EUR, and 146 AMR) using 20, 000 chromosome 21 variants with donor MAF ≥0.01. Five hundred non-admixed synthetic comparator diploids per source population and 2, 000 three-way admixed diploids were projected into this donor-defined space. Extended QC metrics are included in **Supplement Data 1**.

For **Figure 4D**, timing used chr21 admixture-generation runs for 5k, 10k, 25k, and 100k admixed diploids under both the two-way and three-way configurations. Simulation time excluded QC time. Timing data was generated using 32 CPUs and 256 GB of memory.

### Phenotype Generation and GWAS Analysis

For null calibration, we generated continuous phenotypes in 150, 000 EUR and 65, 000 AFR synthetic individuals, with both fixed and background genetic variance components set to zero, and tested variants on chromosomes 1, 3, 8, and 21. Covariate effects were scaled to explain 20% of total phenotypic variance, and residual noise was scaled to explain the remaining 80%. The resulting phenotype was analyzed with REGENIE using the same covariate adjusted GWAS workflow used for the non-null simulations (see below). Null behavior was evaluated using QQ plots and genomic inflation factors.

To generate the example phenotype used in **Figure 5**, FORGEPHAST was used to generate continuous phenotype for chromosomes, 1, 3, 8, and 21 in 100, 000 EUR and 20, 000 AFR synthetic individuals. Causal variants were selected based on whether the MAF ratio between the two populations was less than or equal 8. Candidate sampling used the configured annotation-weighted gene-window mode. Effect sizes were drawn from a Student t distribution with 4 degrees of freedom, scaled by allele frequency in EUR using a transform of 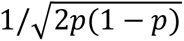, and generated with correlated EUR and AFR effects with target cross-population correlation 0.94 (realized 0.95) and a jitter standard deviation of 0.05. The variance budget assigned 25% of total phenotypic variance to the fixed genetic component, 20% to covariates, and 55% to residual noise, with no background genetic component. Population-stratified GWAS was performed with REGENIE(Mbatchou *et al*., 2021), and recovered effect estimates were compared with the stored causal-variant and effect-size truth tables. Simulation was carried out using 16 CPUs and 128 GB RAM.

GWAS for the null-calibration and phenotype round-trip analyses was performed with REGENIE v4.1(Mbatchou *et al*., 2021) under the quantitative-trait model. For the Figure 5 causal phenotype, EUR and AFR cohorts were analyzed separately. Association models included the simulated non-genetic covariates and global genetic principal components as covariates. Global PCs were computed with PLINK2 after LD pruning using a 200-variant window, 50-variant step size, and *r^2^*=0.2 and the first 100 PCs were included in the REGENIE models. Null calibration used 20 PCs after 1 megabase pruning at *r^2^*=0.1. REGENIE was run using block size 1, 000 in step 1 and block size 400 in step 2. Variant counts, causal-effect summaries, and GWAS recovery metrics are reported in **Supplementary Table 3**.

### Polygenic Risk Score Example

The PRS grid was run on a train/validate/test split of homogenous synthetic AFR/EUR samples, constructed from the same populations as described for **Figure 3** in **Supplementary Table 2**. Only chromosomes 1, 3, 8, and 21 were used as the genotype for this synthetic population, accounting for a total possible causal variant universe of roughly 1.58 million variants. Train was 20K AFR/100K EUR and each of validate and test were 25K of each population, chosen randomly.

54 phenotypes were constructed from a grid across 4 options: heritability (0.05, 0.25, 0.5), polygenicity (0.05%, 0.1%, and 0.5% of the genome as causal), MAF ratio deviation (2-fold, 8-fold, 300-fold), and whether effects were equivalent across populations (jittered as described above/unjittered). The same effect size distribution and MAF scaling used to generate the phenotype in **Figure 5** were used here as well. Only variants with at least 0.05% MAF globally were included. Phenotypic effect size and variance calibration were generated only using train genotypes, with targeted heritability and applied to eval sets. As such, heritability within eval cohorts can differ drastically. Non-genetic covariates and residual variance were also used but were not correlated with ancestry. Exact parameters can be found in available scripts.

GWAS was run separately in the EUR and AFR H2 training strata using REGENIE v4.1 on chromosomes 1, 3, 8, and 21. Per-stratum genotype PCs were precomputed from training genotypes: variants were restricted to SNPs with MAF ≥ 0.05, LD-pruned with PLINK2 *--indep-pairwise 200 50 0.2* and summarized as 100 PCs. These were used as covariates in REGENIE. REGENIE step 1 used the same externally LD-pruned predictor set for whole-genome regression/LOCO prediction (block size 1000), and step 2 tested association across the analysis variants using the step 1 prediction list (block size 400), with the same stratum keep file and PC covariates in both steps.

The 8 PRS tools were then run utilizing the 108 generated summary statistics. Most tools were run either with an explicit grid of options or learnable parameter options. Where tuning was required, configurations were selected separately for each target population by squared Pearson correlation in the matching ancestry-specific validation cohort. Each selected EUR- and AFR-target score was standardized using its matching validation mean and population standard deviation, and the locked transformation was applied to test samples. EUR-target and AFR-target scores were then routed to EUR and AFR test individuals, respectively, before overall, ancestry-specific, and population-centered performance was calculated. Group centering was used only for evaluation and did not affect model fitting or selection. A summary of these grids, as well as custom inputs for train, such as in-sample LD matrices, and any code changes or runtime shims, are summarized in **Supplementary Table 4.**

Final PRS performance was evaluated on the held-out test split as *r^2^*, with overall, ancestry-specific, and population-centered values reported. For the group-centered metric, both phenotype and PRS score were mean-centered within EUR and AFR before computing squared correlation. When normalized performance is shown, *r^2^* was divided by realized test-set *h^2^*, computed in the corresponding evaluation scope as Var(genetic component) / Var(phenotype). Results, as well as GWAS metrics are reported in the **Supplementary Data 2**. Generative AI tools, including OpenAI Codex and Anthropic Claude, were used to assist with code development, documentation, consistency checking, figure preparation, and language editing. Scientific interpretations and final decisions were made by the authors, and all AI-assisted outputs were reviewed and validated by the authors, who take full responsibility for the work.

---

**Supplementary Table 1:** A filled default-threshold cell denotes a hard gate under the default configuration; blank threshold cells denote diagnostics that are reported but not used for default pass/fail decisions.

| Metric | Statistic/ Output | Default threshold | Purpose / interpretation |
| --- | --- | --- | --- |
| Pairwise FST | Between-population pairwise Hudson FST. if the donor baseline is effectively zero, absolute difference is used. | **$\leq 0.20$ relative change**<br>**$\leq 0.30$ relative change for sample mode** | Primary population-structure gate. Synthetic populations should preserve donor-panel differentiation for the same population labels. |
| Within-population split FST | Hudson FST between two subsets drawn from the same synthetic population. | **$\leq 0.002$ absolute FST** | Internal homogeneity gate. Detects artificial substructure within a synthetic population. |
| Stochastic donor-resampling FST | Empirical probability that donor-resampling replicates have mean absolute pairwise FST deviation at least as large as the observed synthetic-vs-donor deviation. | **$\geq 0.05$** | Finite-sample calibration gate. Prevents donor-panel sampling variance from being treated as a simulation failure. |
| LD decay | Weighted RMSE between donor and synthetic median $r^2$ decay curves; relative difference in half-decay distance. AUC of the $r^2$ curve is reported descriptively. | **RMSE $\leq 0.02$; half-decay relative difference $\leq 0.20$** | Local haplotype-correlation gate. Confirms that synthetic LD decay follows donor LD decay across genetic distance; |
| PCA | Difference in population-separation AUC between donor PC space and synthetic samples projected into donor-fit PC space, as well as PC1/PC2 plot | Descriptively reported | Visual and quantitative diagnostic for preservation of donor population structure. PCs are fit on donors and synthetic samples are projected into that space. |
| KING | KING-robust kinship distributions for within-donor, within-synthetic, and cross-synthetic pairs; within-vs-cross synthetic kinship AUC. | Descriptively reported | Diagnostic for individual-level relatedness structure and kinship-based population separation. |
| Allele frequency spectrum | Within-population donor/synthetic MAF histogram distances: total variation, Jensen-Shannon divergence, and Kolmogorov-Smirnov distance. | Descriptively reported | Diagnostic for whether synthetic within-population allele-frequency distributions match donor panels. |
| Runs of homozygosity | Per-individual total ROH length, number of ROH segments, and maximum ROH segment length. | Descriptively reported | Secondary cohort-QC diagnostic for long homozygous tracts and autozygosity-like structure. |
| Hardy-Weinberg equilibrium | Distribution of capped HWE p-values after MAF and count filters | Descriptively reported | Secondary genotype-QC diagnostic. Reported by default |
| Inbreeding F | Distribution of per-variant inbreeding F summaries | Descriptively reported | Secondary genotype-QC diagnostic. Reported by default |

**Supplementary Table 2:**
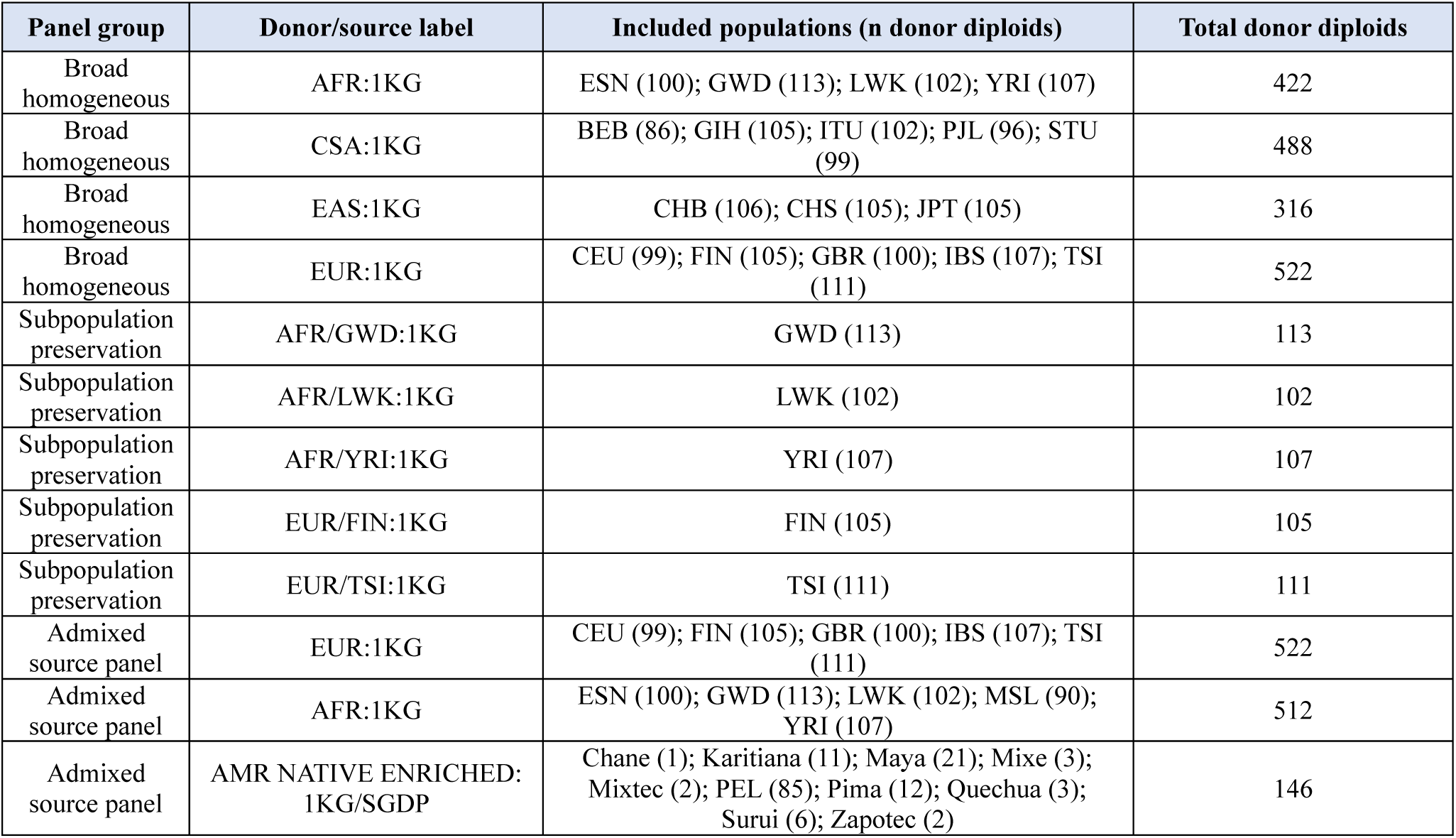
Source populations for various experiments used in both unweighted and weighted mosaic and admixture population generation. 1KG: 1000Genomes (v3), SGDP Simons Genome Diversity Project(Mallick *et al*., 2016)

**Supplementary Table 3:** Figure 5 phenotype-truth and GWAS recovery summary in EUR and AFR cohorts.

|  | EUR | AFR |
| --- | --- | --- |
| Chromosome scope | chromosomes 1, 3, 8, and 21 |  |
| Sample size | 100,000 | 20,000 |
| Target fixed-effect $h^2$ | 0.25 | |
| Realized fixed-effect $h^2$ | 0.229 | 0.296 |
| Pooled realized $h^2$ | 0.251 | |
| Candidate variant universe before tightness filter | 1,579,211 |  |
| Candidate variant universe after tightness filter | 800,285 |  |
| Realized causal variants | 1,579 |  |
| Causal MAF strata (pooled) | 0.05-0.1%: 1; 0.1-0.5%: 14; 0.5-1%: 14; 1-5%: 265; 5-50%: 1,285 |  |
| MAF tightness tier | moderate (8.0-fold cap) |  |
| Effect distribution | Student-t (df=4) |  |
| Base beta SD | 0.15 |  |
| Observed paired EUR-AFR effect correlation | 0.95 |  |
| Effect jitter SD | 0.05 |  |
| GWAS variants tested | 1,248,723 | 1,436,457 |
| Genomic inflation factor | 1.395 | 1.087 |
| Maximum $-\log_{10}(P)$ | 74.8 | 37.8 |
| Genome-wide significant variants ($P < 5e-8$) | 12,818 | 667 |
| Causal variants present in GWAS output | 1,579 |  |
| Beta-recovery Pearson r | 0.932 | 0.852 |

**Supplementary Table 4:**
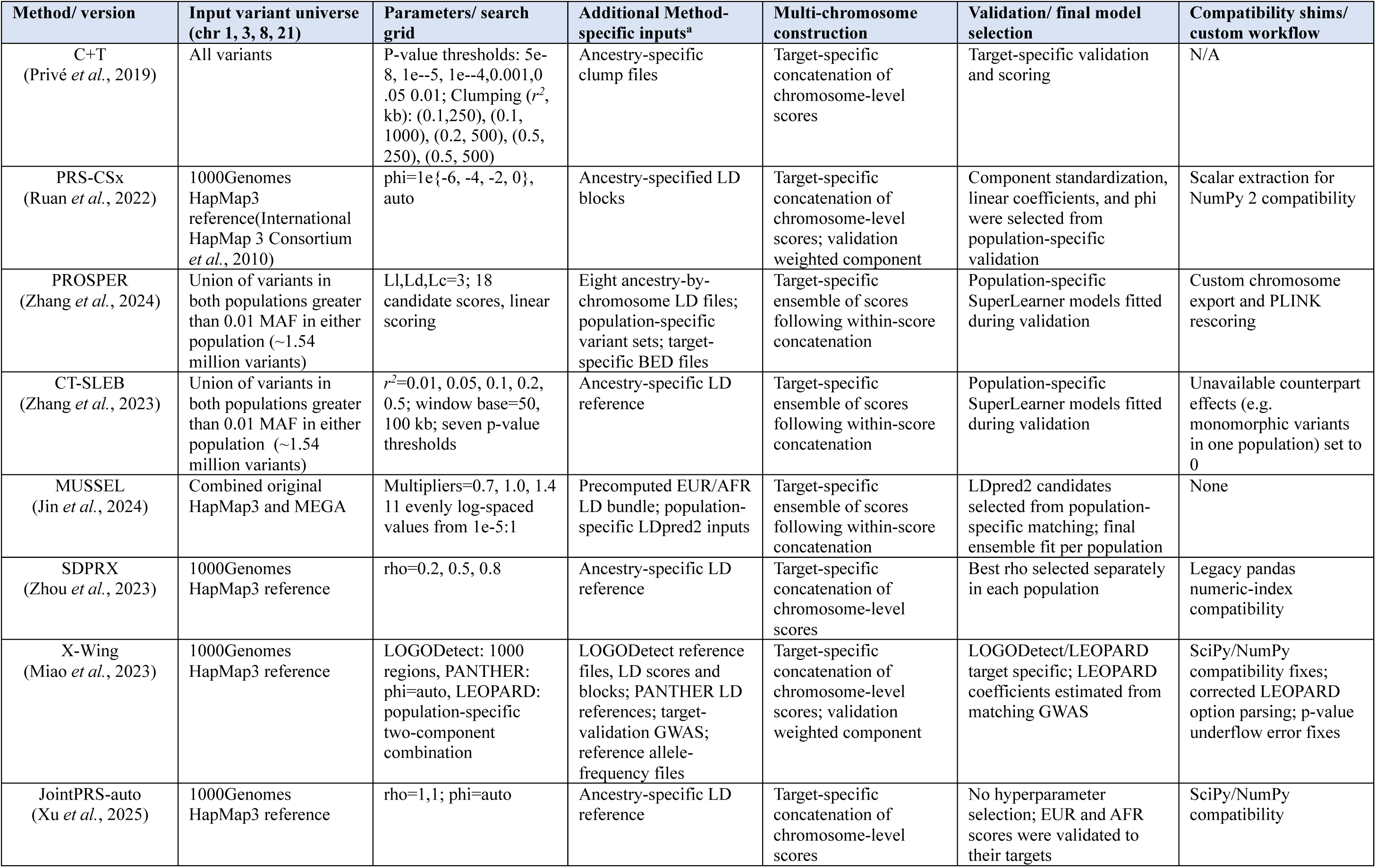
Polygenic risk score method settings used for the FORGEPHAST benchmark. The tab workflow used and any workflow adaptations beyond what is described in the original manuscript/Github for ea chromosome 1, 3, 8, and 21 benchmark. **^a^** All methods received population-stratified GWAS summary statistics

| Method/ version | Input variant universe (chr 1, 3, 8, 21) | Parameters/ search grid | Additional Method-specific inputs<sup>a</sup> | Multi-chromosome construction | Validation/ final model selection | Compatibility shims/ custom workflow |
| --- | --- | --- | --- | --- | --- | --- |
| C+T<br>(Privé *et al.*, 2019) | All variants | P-value thresholds: 5e-8, 1e-5, 1e-4, 0.001, 0.05 0.01; Clumping ($r^2$, kb): (0.1, 250), (0.1, 1000), (0.2, 500), (0.5, 250), (0.5, 500) | Ancestry-specific clump files | Target-specific concatenation of chromosome-level scores | Target-specific validation and scoring | N/A |
| PRS-CSx<br>(Ruan *et al.*, 2022) | 1000Genomes HapMap3 reference (International HapMap 3 Consortium *et al.*, 2010) | $\phi = 1e\{-6, -4, -2, 0\}$, auto | Ancestry-specified LD blocks | Target-specific concatenation of chromosome-level scores; validation weighted component | Component standardization, linear coefficients, and $\phi$ were selected from population-specific validation | Scalar extraction for NumPy 2 compatibility |
| PROSPER<br>(Zhang *et al.*, 2024) | Union of variants in both populations greater than 0.01 MAF in either population (~1.54 million variants) | L1, Ld, Lc=3; 18 candidate scores, linear scoring | Eight ancestry-by-chromosome LD files; population-specific variant sets; target-specific BED files | Target-specific ensemble of scores following within-score concatenation | Population-specific SuperLearner models fitted during validation | Custom chromosome export and PLINK rescoring |
| CT-SLEB<br>(Zhang *et al.*, 2023) | Union of variants in both populations greater than 0.01 MAF in either population (~1.54 million variants) | $r^2=0.01, 0.05, 0.1, 0.2, 0.5$; window base=50, 100 kb; seven p-value thresholds | Ancestry-specific LD reference | Target-specific ensemble of scores following within-score concatenation | Population-specific SuperLearner models fitted during validation | Unavailable counterpart effects (e.g. monomorphic variants in one population) set to 0 |
| MUSSEL<br>(Jin *et al.*, 2024) | Combined original HapMap3 and MEGA | Multipliers=0.7, 1.0, 1.4<br>11 evenly log-spaced values from 1e-5:1 | Precomputed EUR/AFR LD bundle; population-specific LDpred2 inputs | Target-specific ensemble of scores following within-score concatenation | LDpred2 candidates selected from population-specific matching; final ensemble fit per population | None |
| SDPRX<br>(Zhou *et al.*, 2023) | 1000Genomes HapMap3 reference | $\rho=0.2, 0.5, 0.8$ | Ancestry-specific LD reference | Target-specific concatenation of chromosome-level scores | Best $\rho$ selected separately in each population | Legacy pandas numeric-index compatibility |
| X-Wing<br>(Miao *et al.*, 2023) | 1000Genomes HapMap3 reference | LOGODetect: 1000 regions, PANTHER: $\phi$=auto, LEOPARD: population-specific two-component combination | LOGODetect reference files, LD scores and blocks; PANTHER LD references; target-validation GWAS; reference allele-frequency files | Target-specific concatenation of chromosome-level scores; validation weighted component | LOGODetect/LEOPARD target specific; LEOPARD coefficients estimated from matching GWAS | SciPy/NumPy compatibility fixes; corrected LEOPARD option parsing; p-value underflow error fixes |
| JointPRS-auto<br>(Xu *et al.*, 2025) | 1000Genomes HapMap3 reference | $\rho=1, 1$; $\phi$=auto | Ancestry-specific LD reference | Target-specific concatenation of chromosome-level scores | No hyperparameter selection; EUR and AFR scores were validated to their targets | SciPy/NumPy compatibility |

**Supplementary Figure 1.**
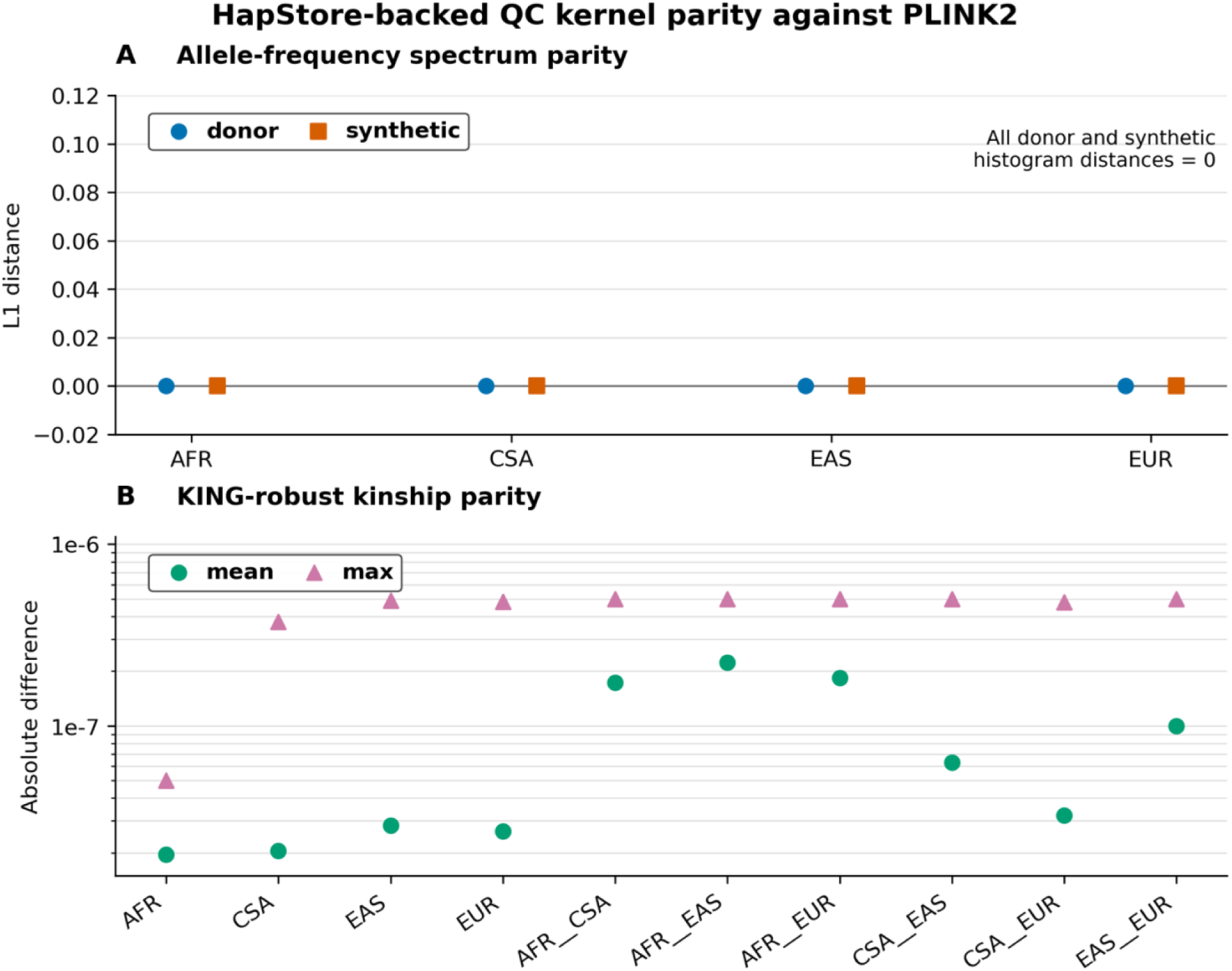
Numerical parity between HapStore-backed FORGEPHAST Python QC kernels and PLINK2 on chromosome 21. **A:** Allele-frequency spectrum histogram distances between HapStore/Python outputs and PLINK2 frequency-count outputs for donor and synthetic cohorts, evaluated at 5, 000 sampled positions using 200 donor and 200 synthetic individuals per population. **B:** Mean and maximum absolute differences between HapStore/Python KING-robust kinship values and PLINK2 KING-table values for 1, 000 sampled pairs per within-population and cross-population comparison.

**Supplementary Figure 2.**
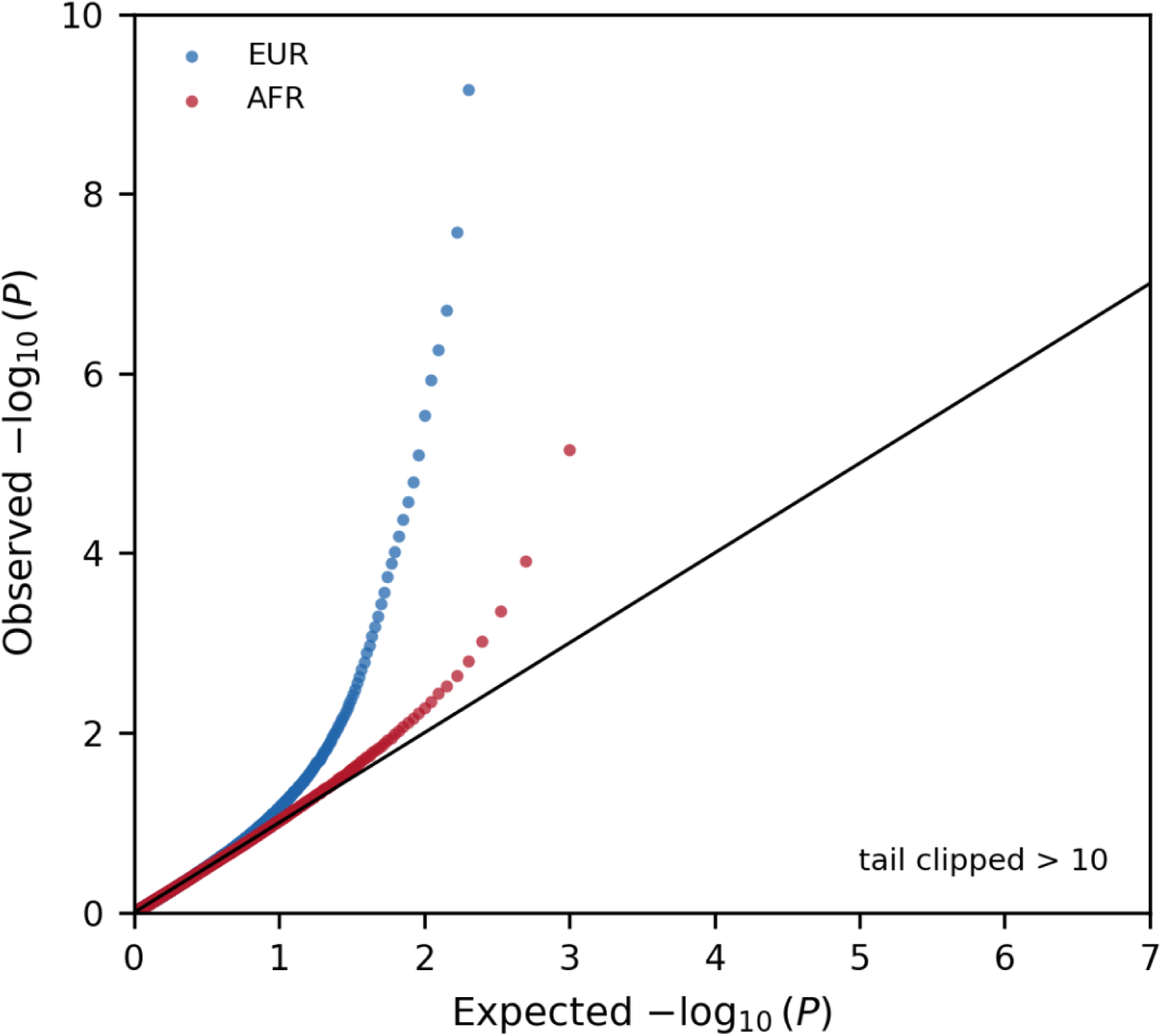
Genomic-control-calibrated quantile-quantile plots for the Figure 5 causal-phenotype genome-wide association analyses. Expected and observed -log10(P) values are shown for the EUR (n = 100, 000) and AFR (n = 20, 000) population-stratified analyses. Association chi-square statistics were divided by the corresponding cohort-specific genomic inflation factors (lambda GC = 1.395 for EUR and 1.087 for AFR) before P values were recalculated (Devlin and Roeder, 1999). Observed -log10(P) values above 10 are clipped for visualization; the black diagonal indicates the expected null distribution

**Supplementary Figure 3.**
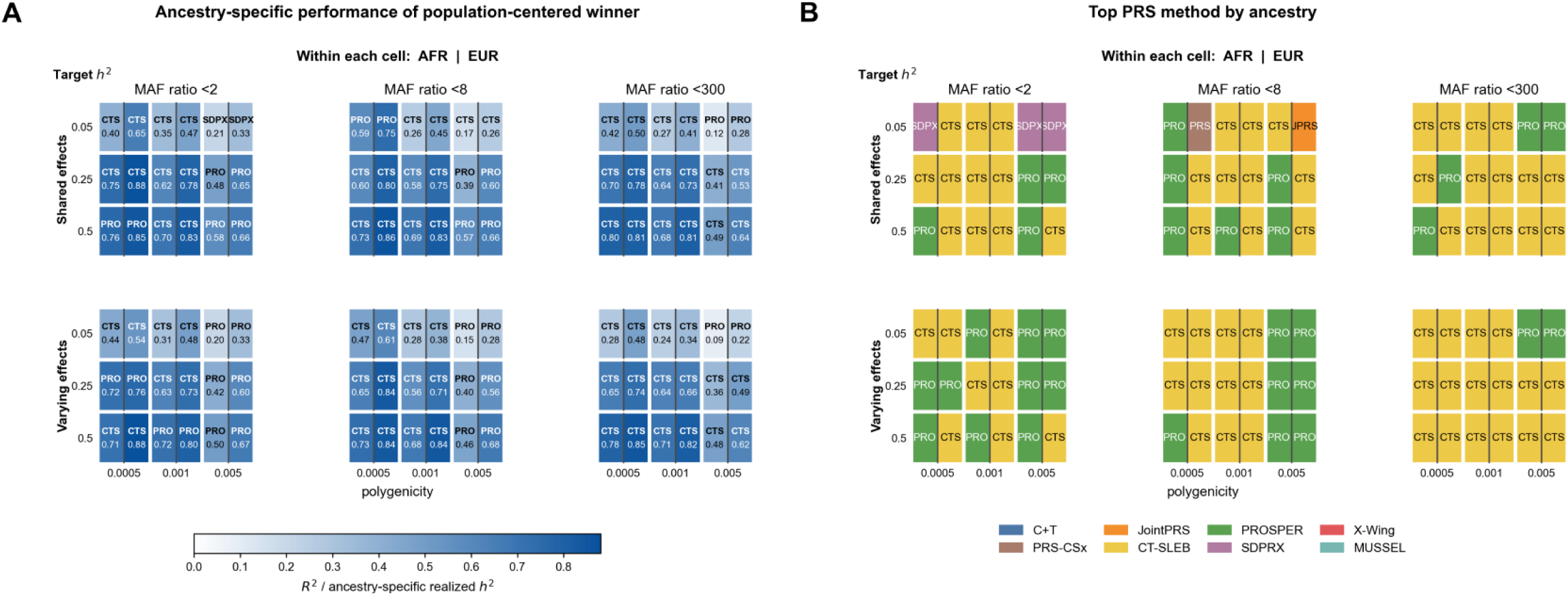
Ancestry-specific polygenic risk score (PRS) performance across 54 quantitative phe Architectures cross target h² (0.05, 0.25, and 0.5), polygenicity (0.0005, 0.001, and 0.005), minor-allele-frequenc and <300), and shared or varying population effect sizes. Within each cell, AFR is shown on the left and EUR on PRS method with the highest population-centered test R² for each phenotype is held fixed; cell values show that specific test R² divided by ancestry-specific realized h². **B:** Colors indicate the top-performing PRS method selec and EUR using ancestry-specific test R² divided by ancestry-specific realized h². Different colors within a cell in ancestry-specific winner

## Works Cited

1000 Genomes Project Consortium et al. (2015) A global reference for human genetic variation. Nature, 526, 68–74.

Abdellaoui, A. et al. (2023) 15 years of GWAS discovery: Realizing the promise. Am. J. Hum. Genet., 110, 179–194.

Abraham, G. et al. (2017) FlashPCA2: principal component analysis of Biobank-scale genotype datasets. Bioinformatics, 33, 2776–2778.

Baumdicker, F. et al. (2022) Efficient ancestry and mutation simulation with msprime 1.0. Genetics, 220.

Chang, C.C. et al. (2015) Second-generation PLINK: rising to the challenge of larger and richer datasets. Gigascience, 4, s13742–015–0047–8.

Corpas, M. et al. (2025) Bridging genomics’ greatest challenge: The diversity gap. Cell Genomics, 5, 100724.

Czech et al. (2025) Analysis-ready VCF at Biobank scale using Zarr. GigaScience, 14.

Falconer, D.S. (1965) The inheritance of liability to certain diseases, estimated from the incidence among relatives. Ann. Hum. Genet., 29, 51–76.

Gower, G. et al. (2025) Accessible, realistic genome simulation with selection using stdpopsim. BioRxiv.

Gravel, S. (2012) Population genetics models of local ancestry. Genetics, 191, 607–619.

Hill, W.G. et al. (2008) Data and theory point to mainly additive genetic variance for complex traits. PLoS Genet., 4, e1000008.

Hou, K. et al. (2024) Admix-kit: an integrated toolkit and pipeline for genetic analyses of admixed populations. Bioinformatics, 40.

Hu, S. et al. (2025) Fine-scale population structure and widespread conservation of genetic effect sizes between human groups across traits. Nat. Genet., 57, 379–389.

Jin, J. et al. (2024) MUSSEL: Enhanced Bayesian polygenic risk prediction leveraging information across multiple ancestry groups. Cell Genomics, 4, 100539.

Kachuri, L. et al. (2024) Principles and methods for transferring polygenic risk scores across global populations. Nat. Rev. Genet., 25, 8–25.

Kalia, S.S. et al. (2024) Development of a breast cancer risk prediction model integrating monogenic, polygenic, and epidemiologic risk. Cancer Epidemiol. Biomarkers Prev., 33, 1490–1499.

Kelleher, J. et al. (2019) Inferring whole-genome histories in large population datasets. Nat. Genet., 51, 1330–1338.

Lauffer, P. et al. (2025) Polygenic risk scores in routine genetic diagnostics: what lies ahead? J. Community Genet., 17, 8.

Liang, M. and Nielsen, R. (2014) The lengths of admixture tracts. Genetics, 197, 953–967.

Li, N. and Stephens, M. (2003) Modeling linkage disequilibrium and identifying recombination hotspots using single-nucleotide polymorphism data. Genetics, 165, 2213–2233.

Mallick, S. et al. (2016) The Simons Genome Diversity Project: 300 genomes from 142 diverse populations. Nature, 538, 201–206.

Manichaikul, A. et al. (2010) Robust relationship inference in genome-wide association studies. Bioinformatics, 26, 2867–2873.

Mbatchou, J. et al. (2021) Computationally efficient whole-genome regression for quantitative and binary traits. Nat. Genet., 53, 1097–1103.

Messer, P.W. (2013) SLiM: simulating evolution with selection and linkage. Genetics, 194, 1037–1039.

Miao, J. et al. (2023) Quantifying portable genetic effects and improving cross-ancestry genetic prediction with GWAS summary statistics. Nat. Commun., 14, 832.

Miles, A. et al. (2020) zarr-developers/zarr-python: v2.4.0. Zenodo.

Nelson, D. et al. (2020) Accounting for long-range correlations in genome-wide simulations of large cohorts. PLoS Genet., 16, e1008619.

Privé, F. et al. (2019) Making the most of clumping and thresholding for polygenic scores. Am. J. Hum. Genet., 105, 1213–1221.

Ralph, P. et al. (2020) Efficiently summarizing relationships in large samples: A general duality between statistics of genealogies and genomes. Genetics, 215, 779–797.

Reich, D. et al. (2009) Reconstructing Indian population history. Nature, 461, 489–494.

Ruan, Y. et al. (2022) Improving polygenic prediction in ancestrally diverse populations. Nat. Genet., 54, 573–580.

Schoech, A.P. et al. (2019) Quantification of frequency-dependent genetic architectures in 25 UK Biobank traits reveals action of negative selection. Nat. Commun., 10, 790.

Sun, G. et al. (2025) HAP-SAMPLE2: data-based resampling for association studies with admixture. Bioinformatics, 41.

Su, Z. et al. (2011) HAPGEN2: simulation of multiple disease SNPs. Bioinformatics, 27, 2304–2305.

Tagami, D. et al. (2024) tstrait: a quantitative trait simulator for ancestral recombination graphs. Bioinformatics, 40.

Tang, Y. and Liu, X. (2019) G2P: a Genome-Wide-Association-Study simulation tool for genotype simulation, phenotype simulation and power evaluation. Bioinformatics, 35, 3852–3854.

Wharrie, S. et al. (2023) HAPNEST: efficient, large-scale generation and evaluation of synthetic datasets for genotypes and phenotypes. Bioinformatics, 39.

Xu, L. et al. (2025) JointPRS: A data-adaptive framework for multi-population genetic risk prediction incorporating genetic correlation. Nat. Commun., 16, 3841.

Zhang, H. et al. (2023) A new method for multiancestry polygenic prediction improves performance across diverse populations. Nat. Genet., 55, 1757–1768.

Zhang, J. et al. (2024) An ensemble penalized regression method for multi-ancestry polygenic risk prediction. Nat. Commun., 15, 3238.

Zhou, G. et al. (2023) SDPRX: A statistical method for cross-population prediction of complex traits. Am. J. Hum. Genet., 110, 13–22.

## Supplementary References

Devlin, B. and Roeder, K. (1999) Genomic control for association studies. Biometrics, 55, 997–1004.

Frankish, A. et al. (2023) GENCODE: reference annotation for the human and mouse genomes in 2023. Nucleic Acids Res., 51, D942–D949.

Hou, K. et al. (2023) Causal effects on complex traits are similar for common variants across segments of different continental ancestries within admixed individuals. Nat. Genet., 55, 549–558.

Hudson, R.R. et al. (1992) Estimation of levels of gene flow from DNA sequence data. Genetics, 132, 583–589.

International HapMap 3 Consortium et al. (2010) Integrating common and rare genetic variation in diverse human populations. Nature, 467, 52–58.

Liang, M. and Nielsen, R. (2014) The lengths of admixture tracts. Genetics, 197, 953–967.

